# Integration of cortical inputs in the lateral hypothalamus is dominated by the medial prefrontal cortex

**DOI:** 10.1101/2024.11.15.623801

**Authors:** L.J.A.M. Razenberg, P.N. de Greef, H.D. Mansvelder, M.M. Karnani

## Abstract

The lateral hypothalamus (LH) is a critical brain region orchestrating survival behaviours including feeding. Its sparse intrinsic synaptic connectivity allows long-range projections to modulate its activity. Some of these projections arise from the cerebral cortex, which is known to influence feeding. However, the functional and anatomical organization of cortico-hypothalamic pathways have remained poorly studied. We used anatomical and optogenetic mapping to show that the medial prefrontal cortex (mPFC) is the strongest cortical input source to the LH, followed by a lateral associative region including the insular cortex (IC), and the ventral subiculum. Input from the mPFC and IC had markedly different synaptic dynamics and were integrated supralinearly. IC input surpassed that of the mPFC in a subpopulation of highly excitable dorsal LH neurons which had a strong h-current. Input from the mPFC showed selective targeting to LH neurons which project back to the mPFC, suggesting the existence of a direct feedback loop. Overall, these results identify a direct prefrontal hypothalamic pathway which is poised to dominate rapid cortical control of hypothalamic activity.

## Introduction

The lateral hypothalamus (LH) is a crucial regulator of arousal, energy balance, and motivated behaviours such as feeding ^1^. Given the sparse local synaptic connectivity within the LH ^2,3^, external inputs can strongly influence the firing rates of LH neurons, particularly at rapid behavioural timescales. One important set of direct inputs arises from the cerebral cortex ^4–6^. However, the functional properties of cortical inputs have remained poorly studied.

The neocortex has a strong influence on feeding. In humans, fronto-temporal lesions often lead to eating disorder symptoms ^7^, and altered cortical volume and functional activity in frontal and insular cortices have been observed in individuals with eating disorders ^8–12^. The medial prefrontal cortex (mPFC) and insular cortex (IC) contain neurons encoding sensory properties of food ^13–15^, some of which are strongly modulated by satiety ^13,15^, consistent with cell-type specific roles in regulating aspects of feeding from food approach to meal duration ^16–23^. Hippocampal volume is also affected in eating disorders ^24^, and hippocampal activity contributes to the control of food intake ^25–28^. These and experimentally induced cortical effects on feeding are likely mediated, at least in part, through direct projections to the LH ^19–22,27–30^, which is a key controller of food consumption ^31^. However, the properties of cortico-hypothalamic pathways are not well-characterized.

Previous anatomical studies have identified direct cortical inputs to the LH ^32–37^, but only few studies have confirmed direct monosynaptic connections using electrophysiological methods ^20,22,29,38^. To understand the mechanisms underlying cortical control of hypothalamic feeding networks, we set out to study the circuit architecture of cortico-hypothalamic pathways using a combination of retrograde tracing and channelrhodopsin assisted circuit mapping ^39^.

We found that cortical output neurons were in the pyramidal layer of the subiculum and in layers 5 and 6 of associative cortices, and had an excitatory effect on LH neurons that was monosynaptic. The mPFC innervated the highest proportion of LH neurons, followed by the IC, ventral subiculum (vSub), ectorhinal cortex (ECT), and dorsal subiculum (dSub). To find out how LH neurons integrate input across time and across cortical pathways, we assessed the effect of stimulus trains and used dual channel optogenetics to probe the mPFC and IC pathways simultaneously ^40^. Unlike the mPFC pathway, the IC pathway exhibited synaptic facilitation and targeted a more defined dorsolateral population within the LH. Interestingly, we observed supralinear integration in a subset of LH neurons that receive input from both of these cortical sources. In addition, a subpopulation of highly excitable dorsal LH neurons preferentially received IC rather than mPFC input, while mPFC input was preferentially targeted at LH neurons which send axons to the PFC.

Our results reveal shared features across cortico-hypothalamic pathways, such as monosynaptic excitation originating from pyramidal neurons, and distinct characteristics like synaptic dynamics and selective targeting of specific LH subpopulations. This lays the groundwork for future studies into control of motivated behaviours, such as feeding, by cortical circuits.

## Results

### Retrograde tracing identifies cortical populations as potential sources of hypothalamic input

To explore which cortical populations project to the LH, we used the retrograde tracer Cholera Toxin Subunit-B (CTb) conjugated to Alexa Fluorophore (647 or 555). We injected small quantities (30 nl) into the LH (see Table 1 for injection coordinates and details), and harvested brains 7-10 days later to assess ipsilateral retrograde labelling (Fig. 1A-B; N = 3). We observed substantial retrograde labelling in several brain regions (Fig. 1 C-L, Fig. S1, Table S1). Our primary focus was on cortical populations, however for comparative purposes, we analysed key subcortical regions known to project to the LH, finding dense labelling in the amygdala (AMY; Fig. 1H), nucleus accumbens (NAc; Fig. 1I) and lateral septum (LS; Fig. 1J).

**Figure 1.**
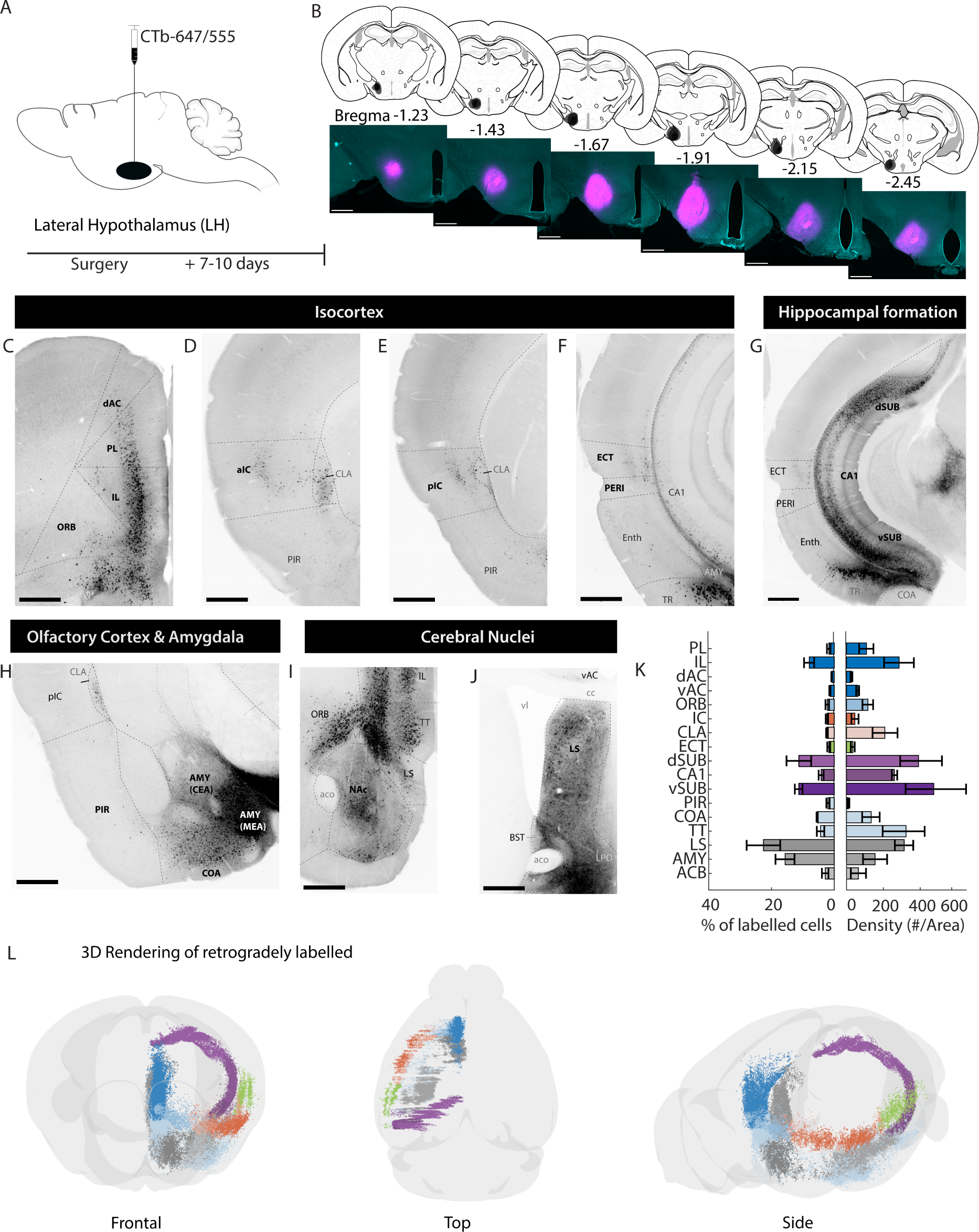
Retrograde tracing from the lateral hypothalamus. **(A)** Schematic of experimental strategy. Cholera toxin Subunit-B (CTb), conjugated to Alexa-647 or 555, was injected into the lateral hypothalamus (LH) and brains were harvested 7-10 days later. See Table 1 for injection coordinates and details. **(B)** Schematic representations of CTb injections into the LH (n = 3 brains), with representative example. Scalebars represent 500µm. **(C – J)** Representative images of retrogradely labelled cells in isocortex, hippocampal formation, amygdala, olfactory cortex, and cerebral nuclei. Scalebars represent 500µm. **(K)** Quantification of retrogradely labelled cells (n=3 brains). Left panel; counts of retrogradely labelled cells per region as a percentage of total of quantified population. Right panel: density of retrogradely labelled cells as counts per area (µm^2^). Data are represented as mean ± SEM (summarized in Table S1). **(L)** 3D views of retrograde labelling (see also supplementary video) Abbreviations follow nomenclature from the Allen Brain Map (mouse.brain-map.org): **AC** (Anterior Cingulate Cortex), **aco** (anterior commissure), **AMY** (Amygdala), **BST** (Bed Nucleus of Stria Terminalis), **CA1** (CA1 Region of the Hippocampus), **cc** (corpus callosum), **CLA** (Claustrum), **COA** (Cortical Amygdala), **ECT** (Ectorhinal Cortex), **Enth** (Entorhinal Cortex), **IC** (Insular Cortex), **IL** (Infralimbic Cortex), **LPO** (Lateral Preoptic Area), **LS** (Lateral Septum), **MEA** (Medial Amygdala), **NAc** (Nucleus Accumbens), **ORB** (Orbitofrontal Cortex), **PERI** (perirhinal cortex), **PIR** (Piriform Cortex), **PL** (Prelimbic Cortex), **SUB** (Subiculum), **TT** (Taenia Tecta), **TR** (Postpiriform transition area), **vl** (Lateral Ventricle). *Directional terms*: **a** (anterior), **p** (posterior), **d** (dorsal) and **v** (ventral).

**Table 1.** Injection coordinates. Anteroposterior (AP), medio-lateral (ML) and dorso-ventral (DV) injection coordinates per experiment and per target area.

| Injectate | Volume (nl) | Target area | AP (mm from Bregma) | ML (mm from Bregma) | DV (mm from brain surface) |
| --- | --- | --- | --- | --- | --- |
| **Experiment 1** |  |  |  |  |  |
| CTb-Alexa647/555 | 30 | LH | -1.3 | 0.95 | -5.4 |
| **Experiment 2** |  |  |  |  |  |
| AAV1-hSyn-ChR2-YFP | 100 | mPFC (unilateral) | 1.8 | 0.4 | -1.9 |
| AAV1-hSyn-ChR2-YFP | 100 | aIC (unilateral) | 1 | 3.2 | -2.7 |
| AAV1-hSyn-ChR2-YFP | 100 total<br>3 * 33 | IC (unilateral) | 1<br>0.5<br>0 | 3.2<br>3.7<br>3.8 | -2.7<br>-2.7<br>-2.8 |
| AAV1-hSyn-ChR2-YFP | 100 | pIC (unilateral) | 0 | 3.8 | -2.8 |
| AAV1-hSyn-ChR2-YFP | 100 | ECT (unilateral) | -3.3 | 4.3 | -2.2 |
| AAV1-hSyn-ChR2-YFP | 100 | dSub (unilateral) | -3.5 | 2.1 | -1.3 |
| AAV1-hSyn-ChR2-YFP | 100 | vSub (unilateral) | -3.8 | 3.3 | -3.2 |
| **Experiment 3** |  |  |  |  |  |
| AAV1-Syn-Chronos-GFP<br>And/or<br>AAV1-Syn-ChrimsonR-tdT | 600 total<br>6 * 100 | mPFC (bilateral) | 1.8<br>1.8<br>2.3 | ± 0.4<br>± 0.4<br>± 0.4 | -1.9<br>-1.5<br>-1.75 |
| AAV1-Syn-Chronos-GFP<br>And/or<br>AAV1-Syn-ChrimsonR-tdT | 600 total<br>6 * 100 | IC (bilateral) | 1<br>0.5<br>0 | ± 3.2<br>± 3.7<br>± 3.8 | -2.7<br>-2.7<br>-2.8 |
| **Experiment 4** |  |  |  |  |  |
| AAV1-hSyn-ChR2-YFP<br>CTb-Alexa647 | 100<br>50 | mPFC (unilateral) | 1.8 | 0.4 | -1.9 |
| AAV1-hSyn-ChR2-YFP<br>CTb-Alexa647 | 100 total<br>3 * 33<br>50 total<br>3 * 17 | IC (unilateral) | 1<br>0.5<br>0 | 3.2<br>3.7<br>3.8 | -2.7<br>-2.7<br>-2.8 |

Within the isocortex, extensive labelling was observed in the mPFC, predominantly in the infralimbic (IL) and prelimbic (PL) areas, with sparser labelling in the anterior cingulate (AC) area (Fig. 1C). As the overall population of labelled neurons in the cortex was continuous across anatomical boundaries and adjacent coronal sections, we generated a 3D rendering from one hemisphere (Fig. 1L). From the mPFC, the labelled population extended ventrally into the olfactory cortex, including the taenia tecta (TT) and dorsal peduncular area (DP; Fig. 1C&I), and caudally into the LS (Fig. 1J).

The labelled population also continued laterally from the IL, DP and TT into orbital areas (ORB; Fig. 1I), extending further into the claustrum (CLA) and the IC (Fig. 1D-E, H), and caudally to the ECT (Fig 1F-G). Labelling in all neocortical regions was primarily restricted to layers 5 and 6, which is comprised mostly of pyramidal neurons (Fig. S1). Additionally, dense labelling was detected in the hippocampal formation, with neurons observed in the pyramidal layer of the dorsal subiculum (dSub), extending ventrally to include the ventral subiculum (vSub) and the CA1 situated between them (Fig. 1G & Fig. S1).

The density of labelling in IL and the PL was comparable to that observed in the LS, suggesting that neocortical inputs to the LH may be as substantial as inputs from traditionally recognized subcortical input hubs like the LS (Fig. 1K).

### Divergent cortical inputs to the LH: mPFC dominance and specialized features of IC inputs

Retrograde tracers are taken up at axons terminating in the injected area but also by (damaged) fibres *en passage*. Retrograde labelling does therefore not guarantee synaptic connectivity. To confirm functional synaptic inputs from the anatomically identified cortico-hypothalamic pathways, we performed channelrhodopsin-assisted circuit mapping (CRACM; ^39^). We injected 100 nl of AAV1-hSyn-ChR2-YFP into one cortical region at a time, allowing for photostimulation of their axon terminals within the LH (Fig. 2A-C, Fig. S2, and Table 1 for injection coordinates and details).

**Figure 2.**
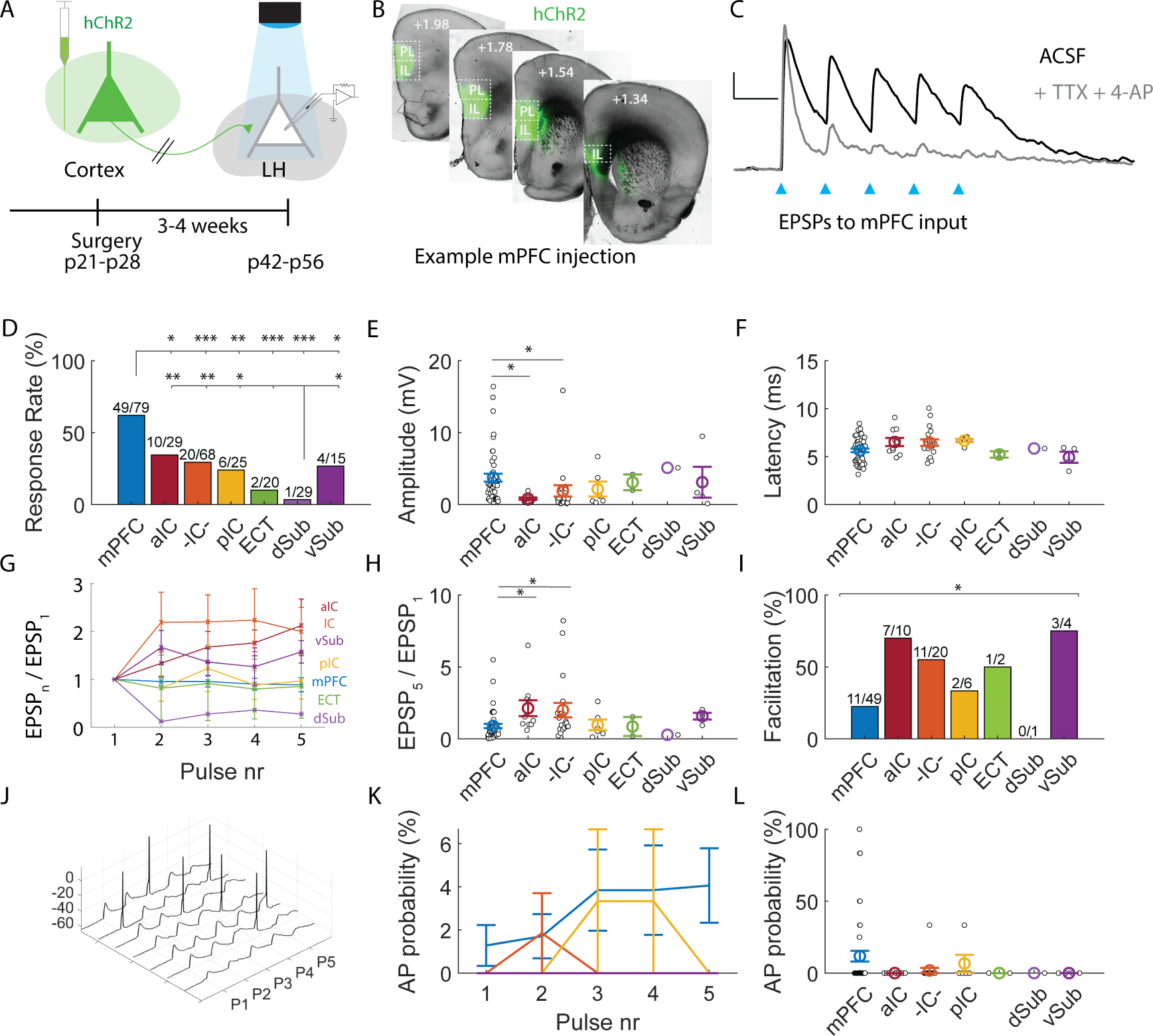
Cortical inputs have distinct synaptic dynamics. **(A)** Schematic and timeline of channelrhodopsin assisted circuit mapping strategy. First, ChR2-yfp was injected into cortex (see Table 1 for injection coordinates and details). After 3-4 weeks time to allow for expression of the virus, brains were harvested for electrophysiology. In ex vivo slices, post-synaptic responses in neurons in the lateral hypothalamic area (LHA) were recorded while neurotransmitter release from ChR2 expressing-cortical axons was triggered with blue light. **(B)** Representative example of viral injection into the medial prefrontal cortex (mPFC), showing 4 different positions from bregma with targeted expression in the infralimbic (IL) and prelimbic (PL) cortex. **(C)** Example of a whole cell-recording (current-clamp) in the LHA. Excitatory post-synaptic responses (EPSPs) were triggered by a train of 5 light pulses (3ms) at 10Hz. Black trace indicates recording in ACSF, while grey trace recording after addition of TTX (1µM) + 4-AP (250µM). Scalebars indicate 3mV and 100ms **(D)** Proportions of LH neurons with postsynaptic responses to photostimulation of cortical axons. The response rate depends on cortical input region (χ^2^ (6, 265) = 46.44, p < 0.0001). A higher proportion of LH neurons is post-synaptic to mPFC compared to the other input, while fewer neurons responded to photostimulation of dSub axons compared to other regions (see Table S2 for pairwise comparisons). **(E)** Response amplitude to the 1^st^ pulse of the stimulus trains differed across cortical input areas Kruskal Wallis χ2 (6, 85) = 18.77, p < 0.01). In particular, initial response amplitudes to mPFC input were larger compared to aIC or -IC- input (U_mPFC-aIC_ = 1362, adj-p = 0.016; U_mPFC-IC_ = 1611, adj-p = 0.025). **(F)** Onset latency of light-evoked EPSPs (pulse 1) in ms. The response latencies were similar across cortical input areas (χ^2^ (6, 84) = 11.47, p = 0.08) **(G)** Summary of paired pulse ratio for each light pulse (n) in the stimulus train (10Hz, 3ms pulses) per input area. A generalized linear mixed effects model revealed significant main effects of pulse number (F (4, 396) = 3,95, p < 0.01), as well as an interaction of pulse number and input area (F (24, 396) = 1.65, p = 0.030). **(H)** The paired pulse ratio of the first and final light pulse (EPSP_5_/EPSP_1_) in the stimulus train (10Hz, 3ms pulses) differed between input areas (χ^2^ (6, 85) = 20.61, p < 0.01). EPSPs to activation of aIC and IC showed more facilitation compared to mPFC input (U_mPFC-aIC_ = 1063, adj-p = 0.014; U_mPFC-IC_ = 1207, adj-p = 0.017). **(I)** Summary of facilitation rate in %, defined as the proportion of neurons with a paired pulse ratio (EPSP_5_/EPSP_1_) larger than 1 (χ^2^ (6, 92) = 14.94, p = 0.021). Pairwise comparisons are non-significant after correction for multiple testing. **(J)** Example of action potential firing in response to photostimulation of mPFC axons. **(K)** Summary of the probability of action potential firing for each light pulse in the stimulus train, per input area. Generalized linear mixed effect model revealed no significant effects of cortex, pulse number or their interaction on spike probability (F_Cortex_ (6, 355) = 0.82, p = 0.552; F_PulseNr_ (4, 355) = 1.99, p = 0.096; F_Cortex*PulseNr_ (24, 355) = 0.82, p = 0.714), but notably action potentials were only observed during mPFC or IC stimulation. **(L)** Summary of the probability of firing at least one action potential during the stimulus train (χ^2^ (6, 77) = 5.54, p = 0.476).

We recorded excitatory postsynaptic potentials (EPSPs) in response to cortical inputs from the mPFC, IC, ECT, dSub, and vSub in LH neurons (Fig. S2, Fig. S3). The IC extends along the rostrocaudal axis and is typically divided into anterior (aIC) and posterior (pIC) regions, each with distinct projection patterns ^41^. Based on previous tracing results (Fig. 1 & Fig. S1), we targeted caudal aIC and rostral pIC with separate injections and performed an additional multi-injection across the rostrocaudal extent to encompass the full projection population (-IC-; Fig S2B-E).

To assess whether responses were monosynaptic, we applied tetrodotoxin (TTX) and 4-aminopyridine (4-AP)^42^. Any optically evoked EPSP persisting in the presence of TTX and 4-AP is attributed to direct activation of ChR2 at the presynaptic terminal, rather than disynaptic input. All tested postsynaptic responses persisted with TTX and 4-AP (mPFC: n=9/9; IC: n=6/6; vSub: n=4/4; Fig. 2C, S2). Pooled responses in TTX+4-AP were 62% larger than in baseline (EPSP_bsl_ = 3.94 ± 1.10 mV vs. EPSP_TTX+4-AP_ = 6.40 ± 1.56 mV; Z = -3.24, p < 0.01). Although TTX+4-AP typically reduces response efficacy ^42,43^, some studies have also reported increases under similar conditions in other circuits ^44–46^. Onset latencies of all recorded EPSPs were consistent with latencies of TTX+4-AP confirmed monosynaptic inputs (Latency_unconfirmed_ = 6.00 ± 0.16 ms vs. Latency_confirmed_ = 5.79 ± 0.30 ms; Z = 0.22, p = 0.829; Fig. 2F). Together, these data indicate that the LH receives monosynaptic input from mPFC, IC, ECT, dSub and vSub.

A significantly higher proportion of LH neurons (n=49/79) responded to inputs from the mPFC than any of the other areas (Fig. 2D and Fig. S2). Inputs from ECT, dSub were the sparsest (Fig. 2D and Fig. S2E&F), despite extensive retrograde labelling in the latter (Fig. 1). Anterograde tracing showed that dSub axons were primarily localized in the fornix, consistent with potential retrograde tracer take-up by fibres of passage (Fig. S3D).

Stimulation of mPFC terminals resulted in larger EPSP amplitudes to the first pulse of 10Hz trains compared to the IC (U_mPFC-IC_ = 1611, adj-p = 0.025) and specifically the aIC (U_mPFC-aIC_ = 1362, adj-p = 0.016, Fig. 2E). However, while the amplitude to the first EPSP was larger for mPFC input, no difference between regions was observed when considering the maximum EPSP amplitude from the train (χ2 (6, n = 85) = 10.54, p = 0.104; not shown). No differences were observed in EPSP latency across cortical injections (Fig. 2F; χ^2^ (6, n = 84) = 11.47, p = 0.075).

Responses to these inputs displayed distinct synaptic dynamics (Fig. 2G). The paired-pulse ratio (EPSP5/EPSP1) was higher for IC and aIC inputs compared to mPFC inputs, indicating facilitation of responses to insular inputs (U_mPFC-aIC_ = 1063, adj-p = 0.014; U_mPFC-IC_ = 1207, adj-p = 0.017; Fig. 2H-I). Action potential (AP) firing was only observed in response to mPFC and IC stimulation (Fig. 2J-L). Anterograde tracing revealed that axonal distributions differ across regions and align with the locations of postsynaptic neurons (Fig. S3). Axons from the mPFC and IC were dense, while ECT, dSub, and vSub axons were sparser in the LH, consistent with the observed response rates (Fig. S3, Fig. 2D). mPFC and IC axons were concentrated in the lateral regions of the hypothalamus, largely avoiding adjacent structures like the ventromedial and dorsomedial nuclei (Fig. S4A-D). However, mPFC axons occupied a larger overall area compared to IC axons (F (1, 20) = 57.91, p < 0.001), which were more restricted to a specialized domain located dorsally (F (1, 20) = 5.40, p = 0.031) and laterally (F (1, 20) = 25.74, p < 0.001) relative to mPFC fibres (Fig. S4E). These findings suggest that different cortical regions contribute uniquely to LH innervation.

### A subpopulation of LH neurons receives convergent mPFC and IC inputs and integrates them supralinearly

Given the differences in anatomical organization between mPFC and IC fibres and their synaptic dynamics, we next asked if they innervate distinct LH neurons. To investigate this, we used a dual-channel optogenetics strategy where each cortical area expressed either the blue-shifted opsin Chronos or the red-shifted ChrimsonR, which can be independently activated by separate light wavelengths ^40^. Chronos is activated by blue (470 nm) light, while ChrimsonR responds to red (590 nm) light but can also be excited by 470 nm light at higher intensities. Therefore, it is crucial to establish a 470 nm light power range that reliably excites Chronos but not ChrimsonR.

To validate this approach, we first injected ChrimsonR and Chronos into separate animals (Fig. S5A, see Table 1 for injection coordinates and details). We recorded AP firing in ChrimsonR- or Chronos-expressing neurons at the injection site in response to 470 nm and 590 nm stimulation across a range of light intensities. As expected, 590 nm light stimulation evoked APs in ChrimsonR-expressing neurons but not in those expressing Chronos (Fig. S5B-D). However, the 470 nm irradiance required to achieve 95-100% firing probability in Chronos-expressing neurons also induced APs in ChrimsonR expressing neurons (Fig. S5B & E), as well as EPSCs in LH neurons in animals only expressing ChrimsonR (Fig. S5F-J). Despite some Chronos-expressing neurons firing at lower light intensities (Fig. S5C-E), the selective activation range without triggering ChrimsonR was suboptimal, with a firing probability of only 83% (± 16% SEM). Notably, lower intensity 470 nm light still induced small EPSCs in the LH of ChrimsonR animals with up to 25% probability. Therefore, we used a high 470 nm irradiance and sequential photostimulation to desensitize ChrimsonR with a 250 ms 590 nm pulse prior to 470 nm light stimulation (Fig. S5K-L) ^47,48^.

We then injected ChrimsonR and Chronos into the mPFC and IC, reversing the injection sites in separate experiments to control for opsin-related effects on synaptic dynamics (Fig. 3A). Bilateral injections were made at multiple locations in the mPFC and IC to ensure coverage of the projection populations. Using the desensitization strategy, ChrimsonR-mediated responses to blue light were suppressed while preserving the postsynaptic responses to Chronos, enabling categorization of LH neurons as responsive exclusively to ChrimsonR, Chronos, both opsins, or neither (Fig. 3B-D).

**Figure 3.**
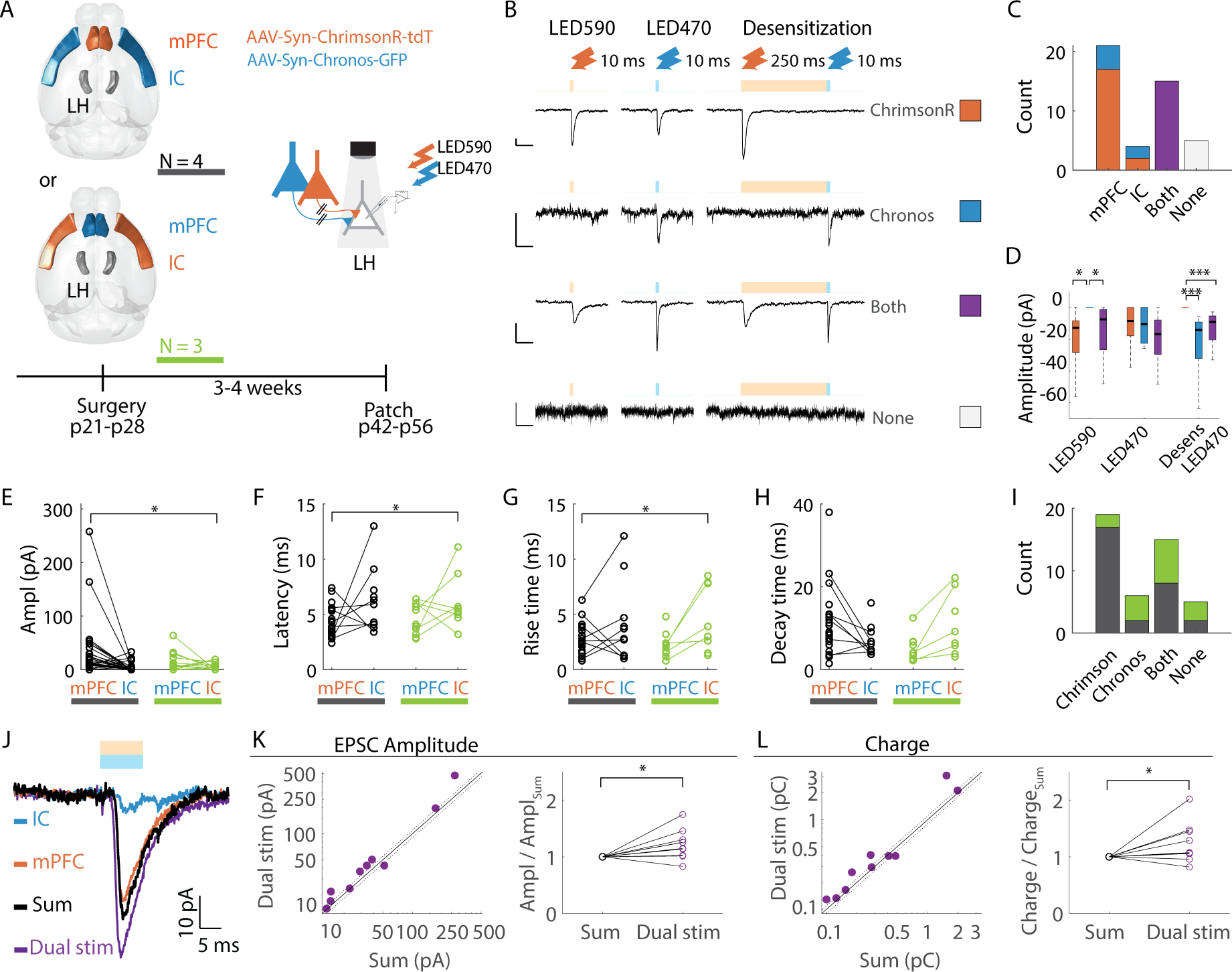
mPFC and IC inputs converge supra-linearly in LH. **(A)** Schematic and timeline of dual channel optogenetics experiment (see Table 1 for injection coordinates and details). Mice are injected with both AAV1-Syn-ChrimsonR-tdT and AAV1-Syn-Chronos-GFP bilaterally into the mPFC and the IC (mPFC_ChrimsonR_ and IC_Chronos,_ N = 4), or vice versa (mPFC_Chronos_ and IC_ChrimsonR,_ N = 3) to account for opsin effects. After expression of virus, whole-cell patch clamp recordings are performed in LH neurons to record postsynaptic responses to photostimulation of ChrimsonR and Chronos terminals. **(B)** Example traces of different postsynaptic responses (top to bottom): response exclusively to ChrimsonR (with 470nm crosstalk suppressed by desensitization), response exclusively to Chronos, response to both **(C)** Cell count of neurons with responses to exclusively mPFC input, exclusively IC input, mPFC and IC input (both), and of neurons with no EPSC to either input. Note that given the two different opsin/area combinations, mPFC and IC responding neurons are made of a combination of ChrimsonR and Chronos responsive neurons. **(D)** Summary of EPSC responses per opsin (ChrimsonR in orange, Chronos in blue, and both in purple) to protocols seen in (B): 10ms stimulation with 590nm light (LED590), 470nm light (LED470) and 470nm light preceded by a desensitizing pulse (Desens LED470). Chronos neurons are not responsive to LED590 (Z_ChrimsonR vs. Chronos_ = -2.91, p = 0.011; Z_Chronos vs. Both_ = 2.35, p = 0.042). All categories responded to LED470, but a preceding desensitizing LED590 pulse suppressed this response exclusively in ChrimsonR neurons (Z_ChrimsonR vs. Chronos_ = 3.88, p < 0.001; Z_ChrimsonR vs. Both_ = 4.66, p < 0.001). **(E-H)** Characteristics of mPFC and IC responses, grouped by opsin injection combination (mPFC_ChrimsonR_ and IC_Chronos_ in black; or mPFC_Chronos_ and IC_ChrimsonR_ in green) to control for opsin effects on EPSC amplitude (**E**, absolute values), latency **(F)**, rise time **(G)**, and decay time **(H)**. Linear mixed-effects analysis reveals significant effects of cortical area, but not opsin, on amplitude (F(1, 86) = 4.02, p = 0.048), latency (F(1, 51) = 7.13, p = 0.010) and rise time (F(1, 49) = 5.07, p = 0.029). **(I)** Absolute cell counts for opsin response categories, showing contributions per cortical injection area (mPFC_ChrimsonR_ and IC_Chronos_ in black; or mPFC_Chronos_ and IC_ChrimsonR_ in green). **(J)** Example traces of LH neuron integrating cortical responses. Recordings of IC (blue trace) and mPFC (orange trace) induced oEPSCs in response to separate 10ms stimulation of 470nm light (desensitized) or 590nm, respectively. The arithmetic sum of the mPFC and IC response (black trace; Sum) is smaller than the response to simultaneous dual-color stimulation (purple trace; dual stim). **(K)** The EPSC amplitudes from the arithmetic sum of mPFC and IC EPSCs are compared against those from simultaneous dual-color stimulation (n=10 cells). The black line represents the linear relationship (where the sum equals dual stimulation), and the dotted lines indicates the 10% confidence intervals. The right panel shows amplitude ratios. Amplitudes are divided by the amplitude of the arithmetic sum to assess linearity. The ratio for dual stimulation is larger than 1 (Z = -2.29, p = 0.022, n=10). **(L)** Same as K, but for the integral of the EPSC, which represents total synaptic charge transfer (pC). The ratio for dual stimulation is larger than 1 (Z = -2.09, p = 0.037, n=10).

Most neurons received input from the mPFC, either exclusively or in combination with IC inputs (n=36/45; 80%). 33.3% of neurons (15/45) integrated inputs from both mPFC and IC, while 46.7% (21/45) received input exclusively from the mPFC, and 8.9% (4/45) received input exclusively from the IC. The occurrence of cells with both mPFC and IC inputs did not significantly differ from a chance overlap (χ² (3, n = 45) = 0.02, p = 0.880), consistent with independent targeting by mPFC and IC inputs.

We compared EPSC parameters using generalized linear mixed-effects models, controlling for the effects of opsin type (Chronos vs. ChrimsonR), injection site (mPFC vs. IC), and their interaction (Fig. 3E-H). EPSCs evoked by mPFC stimulation exhibited significantly larger amplitudes (Fig. 4E; F(1, 86) = 4.02, p = 0.048), shorter latencies (Fig. 3F; F(1, 51) = 7.13, p = 0.010), and faster rise times (Fig. 3G; F(1, 49) = 4.07, p=0.029) compared to those evoked by IC stimulation. No significant effects of opsin type, injection site, or their interaction were observed on EPSC decay times (Fig. 3H).

**Figure 4.**
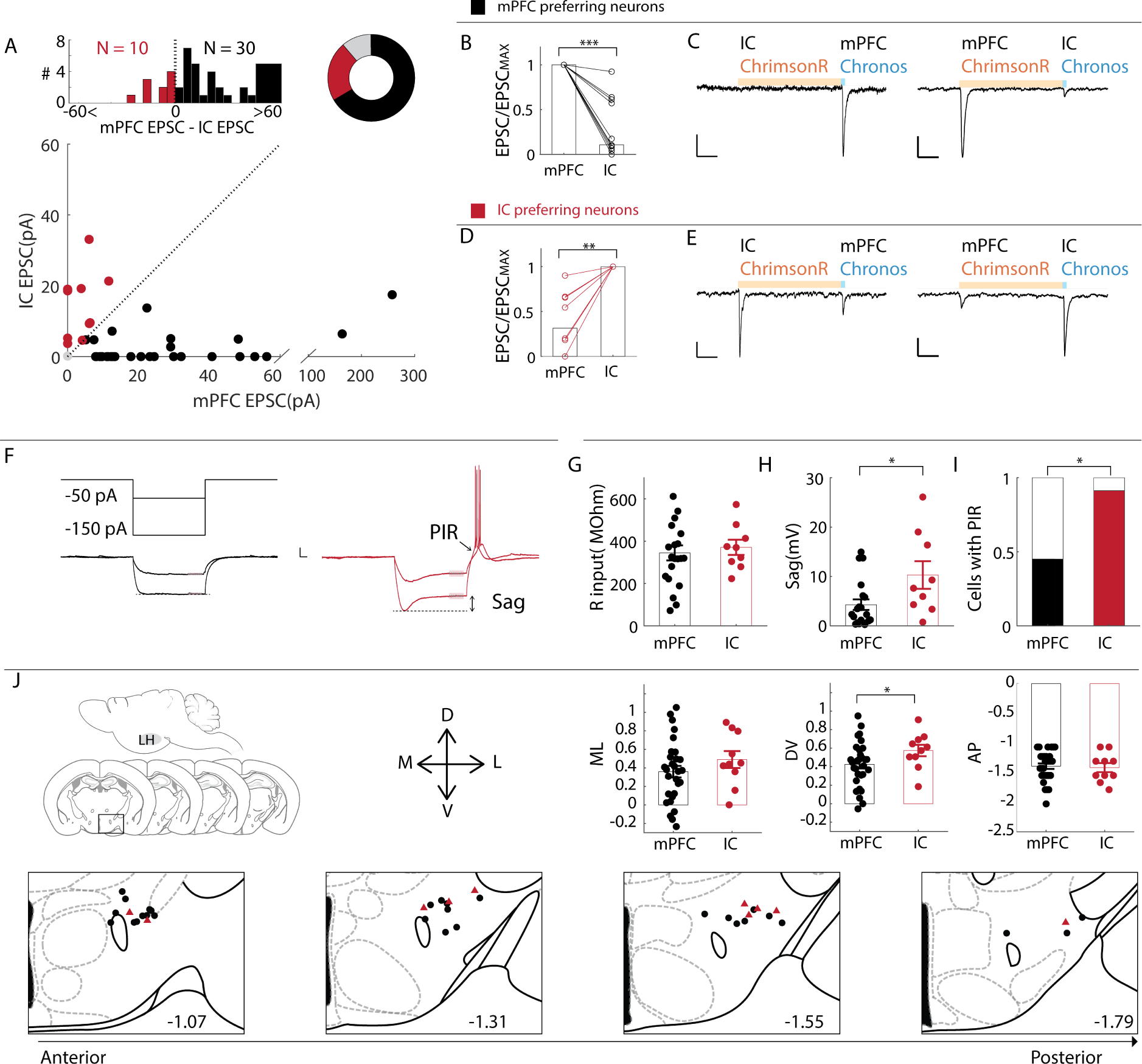
Increased excitability of IC-input preferring dorsal LH neurons. **(A) Bottom panel**: amplitudes of EPSCs in LH neurons to evoked mPFC and IC input in dual-channel optogenetics experiment (see figure 4). The dashed line represents a 1:1 relationship, where the amplitude of the mPFC EPSC is equal to the amplitude of the IC EPSC. **Top panel (left)**: histogram illustrating the distribution of the difference between mPFC – and IC EPSC amplitudes. The difference was computed by subtracted EPSC_IC_ from EPSC_mPFC_. Neurons with larger mPFC EPSC compared to IC EPSC amplitudes (black, n=30) are considered ‘mPFC preferring’, while neurons with larger IC EPSCs compared to mPFC EPSCs are considered ‘IC preferring’ (red, n=10). **Top panel (right):** Pie chart summarizing the total of 45 neurons recorded. mPFC dominated neurons made up the majority of the population (∼67%; n=30/45), followed by IC dominated neurons (∼22%; n=10/45) and a small proportion (∼11%, n=4/45) without detectable mPFC or IC input (grey segment). **(B)** EPSC currents normalized to peak response. mPFC preferring neurons have larger EPSC amplitudes to mPFC compared to IC input (Z = 4.99, p < 0.0001) **(C)** Example traces of LH neurons predominantly receiving mPFC input using two different combinations of opsin. ChrimsonR is stimulated by a 250ms pulse of red light (590nm), followed by a 10ms pulse of blue light (470nm) to stimulate Chronos but not ChrimsonR (see fig 4). Scalebars correspond to 10mV, 10ms. **(D-E)** Same as (B-C) but for LH neurons predominantly receiving IC input, n = 10. IC preferring neurons have larger EPSC amplitudes to IC compared to mPFC input (Z = -2.82, p < 0.01). **(F)** Example traces to negative current steps (-150pA and -50 pA), for a mPFC dominated neuron (black traces) and an IC dominated neuron (red traces). Scalebar corresponds to 10mv, 50ms. Input resistance **(G**; Z = -0.45, p = 0.651) was similar for both input types, but predominantly IC responsive neurons had larger sag voltages **(H**; Z = -2.24, p = 0.025), and more often displayed a post inhibitory rebound (PIR) **(I**; χ^2^ (1, 28) = 5.32, p = 0.021) **(J)** Locations of recorded neurons within the LH. IC preferring neurons were located more dorsally in the LH compared to mPFC preferring neurons (Z=-1.98, p = 0.047). Their mediolateral (Z=-0.80, p=0.426), and anteroposterior (Z = 0.48, p = 0.634) distributions were similar.

In neurons receiving convergent inputs from both mPFC and IC, we examined whether these are integrated linearly. To do this, we compared the summed separate responses to mPFC and IC stimulation to the response to simultaneous dual-colour stimulation (Fig. 3J). If the inputs are integrated linearly, the response to simultaneous stimulation should equal the sum of the separate responses, resulting in a ratio (Dual stimulation / Sum) of 1 while supralinear integration would yield a ratio greater than 1 ^49^. Simultaneous dual-colour stimulation produced larger response amplitudes and greater synaptic charge transfer than the calculated sum (Z_amplitude_ = -2.29, p_amplitude_ = 0.022; Z_charge_ = -2.09, p_charge_ = 0.037, n =10), and their ratios were larger than 1 (Fig 3K-L). These data indicate supralinear summation of mPFC and IC input in LH neurons.

### Neurons preferentially responsive to IC input are highly excitable and clustered in dorsal LH

While, across all our recordings, mPFC input was on average stronger than IC input (Fig. 4A-C), a subset of neurons had stronger IC input (Fig. 4A, D-E), and only a small subset did not receive input from either source. We asked if these distinctly sensitive neurons represent an electrophysiologically distinguishable subtype ^50^. We therefore categorized neurons based on their dominant input type (mPFC-preferring or IC-preferring) and compared their intrinsic properties (Fig. 4F-I & Fig. S6).

Both types displayed similar input resistances (Fig. 4F-G), but IC-preferring neurons showed a more pronounced voltage sag (Z = 2.24, p = 0.025: Fig. 4F, H) in response to hyperpolarizing current steps and a higher incidence of post-inhibitory rebound depolarization (χ^2^ (1, 28) = 5.32, p = 0.021; PIR; Fig. 4H, I). Current ramp measurements revealed no differences in rheobase current, gain, or overall firing rate (Fig. S6A-D), indicating that the observed excitability traits (voltage sag, PIR) are selectively engaged under specific conditions Consistent with IC axon organization (Fig. S4), IC-preferring neurons were located more dorsally within the LH (stats; Fig. 4N).

To further characterize these neurons, we analyzed spontaneous EPSCs (sEPSCs) to explore whether baseline excitatory drive might reveal additional differences in information processing. In cortical regions, neurons with Ih-mediated sag voltage are associated with distinct input profiles, similar to the input preferences observed here, and heightened excitatory synaptic activity ^51–53^. However, we observed that IC-preferring and mPFC-preferring neurons had comparable amplitudes and frequencies of sEPSCs (Fig. S6E-H), suggesting that cortical input sensitivity and voltage sag are independent of excitatory synaptic activity.

To evaluate how cortical inputs contribute to synaptic drive, we compared sEPSCs with optogenetically evoked EPSCs (oEPSCs). Since optogenetic stimulation can potentially activate many synapses simultaneously, while sEPSCs typically reflect individual synaptic events, comparing sEPSCs with optogenetically evoked EPSCs (oEPSCs) provides an estimate of synaptic density or sparsity^2^. We fit cumulative distributions of sEPSCs to compare with oEPSC data and observed that while mPFC oEPSCs were on average larger than 50% of sEPSCs, only 35% of IC oEPSCs exceeded sEPSC amplitude. There was no significant difference in the proportion of sEPSCs smaller than the oEPSCs between both neuron types (Fig. S6H). The similarity in amplitude between oEPSCs and sEPSCs is consistent with a sparse synaptic distribution, suggesting that, although cortical inputs in the LH are abundant, they likely involve only a limited number of synapses. Notably, there was considerable variability in oEPSC amplitude and relative strength across neurons, potentially reflecting cell-specific differences in cortical input strength, which could be modulated by learning processes ^6,20,54^.

Together, these findings indicate that LH neurons are distinctly sensitive to cortical inputs, integrating them based on a preference independent of overall excitatory drive but reflected in their intrinsic properties: IC-preferring neurons in the dorsal LH exhibit increased excitability due a stronger voltage sag and PIR, which can lead to burst firing output ^55^.

### mPFC, but not IC, projections preferentially target reciprocally connected LH neurons

The LH projects to various downstream and upstream targets, and previous tracing work has suggested the LH is connected reciprocally with most of these targets ^34,56,57^. We therefore asked whether mPFC or IC inputs are biased toward reciprocally connected neurons. To investigate this, we co-injected ChR2 and CTb into either the mPFC or the IC (Fig. 5A, see Table 1 for injection coordinates and details).

**Figure 5.**
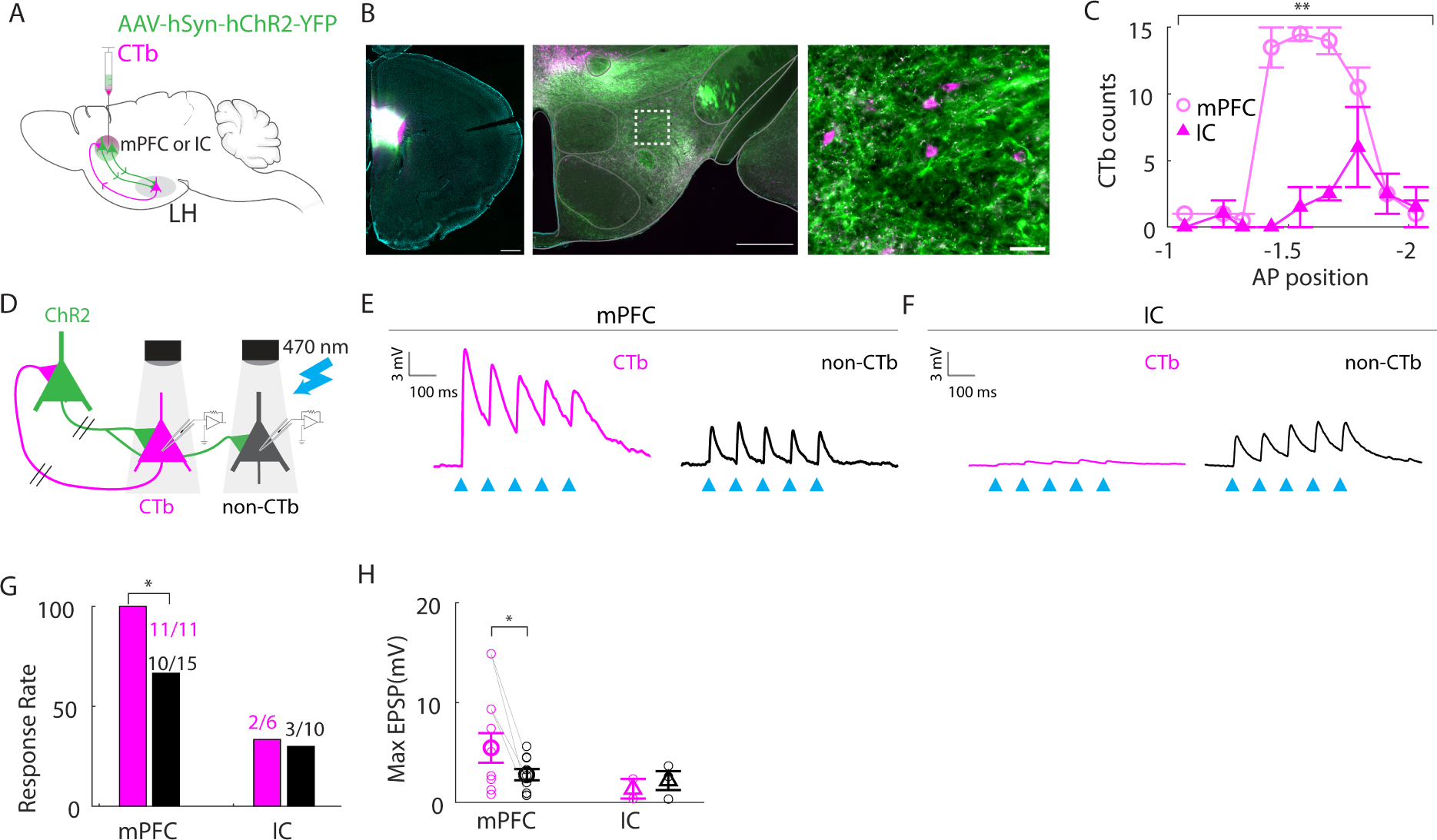
mPFC, but not IC, afferents preferentially target reciprocally connected LHA neurons. **(A)** Schematic of combined injection of AAV-hSyn-hChr2-YFP and retrograde tracer CTb-647 into the mPFC or the IC, to study reciprocal connectivity with the LH. See Table 1 for injection coordinates and details. **(B)** Histological sections. Left panel: example of an injection into the mPFC, showing overlay of CTb-647 and ChR2-YFP in the mPFC (scalebar = 500 µm). Middle panel: ChR2-YFP expressing afferents and retrogradely labelled cell bodies in the LH (scalebar = 500µm). Right panel: zoomed in (scalebar = 50µm). **(C)** Cell counts of CTb positive neurons in LH sections ranging from anterior to posterior, for mPFC injections (n=2) and IC injections (n=2). Generalized Linear Mixed Effects (GLME) model indicates a significant effect of injection site, but not AP-position (F_mPFC-IC_ (1,32) = 11.28, p = 0.002; F_AP-pos_ (1,32) = 2.02, p = 0.002; F_mPFC-IC * AP-pos_ (1,32) = 5.86, p = 0.021). **(D)** Schematic of electrophysiological recording strategy. CTb-positive and -negative neurons in the LH are recorded during photostimulation with 470nm light (5 pulses of 3ms @ 10Hz). **(E)** Example traces of CTb-positive (reciprocal) and -negative (non-reciprocal) neurons in the LH responding to mPFC input with EPSPs. **(F)** Same as E, but for IC. **(G)** Response rate for mPFC and IC injections. The proportion of neurons postsynaptic to mPFC is higher for reciprocally connected neurons compared to non-reciprocal neurons (χ^2^ (1,26) = 4.54, p = 0.033). The proportions of neurons postsynaptic to the IC are similar (χ^2^ (1,16) = 0.019, p = 0.889). **(H)** EPSP amplitudes of CTb^+^ and CTb^-^ neurons responding to mPFC or IC input. Lines indicate neurons recorded simultaneously in a multiple-patch configuration. CTb-positive (reciprocal) neurons have larger responses to mPFC input compared to CTb-negative (non-reciprocal) neurons ( F (1,15) = 6.95, p = 0.019).

Following injections, retrogradely labelled neurons were observed in the LH (Fig. 5B), with a considerable number identified after mPFC injections (n=2 mice), particularly in intermediate anterior-posterior positions (-1.43 to -1.79 mm from bregma; Fig. 5C). In contrast, IC injections (n=2 mice) resulted in a significantly lower number of retrogradely labelled neurons in the LH (GLME model; F_mPFC-IC_ (1,32) = 11.28, p < 0.01; F_AP-pos_ (1,32) = 2.02, p < 0.01; F_mPFC-IC * AP-pos_ (1,32) = 5.86, p = 0.021).

We targeted retrogradely labelled neurons to assess their responsiveness to reciprocal cortical input. 100% (10/10) of LH->mPFC neurons received mPFC input, compared to 67% of non-mPFC projecting neurons (10/15), a significant difference (χ^2^ (1,26) = 4.54, p = 0.033). In contrast, IC axon stimulation elicited similar response rates in IC- and non-IC projecting LH neurons (33% (2/6) vs. 30% (3/10); χ^2^ (1,16) = 0.02, p = 0.889, Fig. 5G). Moreover, mPFC-projecting LH neurons had larger EPSP amplitudes in response to mPFC input compared to non-mPFC projecting LH neurons (F (1,15) = 6.95, p = 0.019). We observed no differences in EPSP latency (CTb+ = 5.19 ± 0.43; CTb- = 5.16 ± 0.40; F (1,15) = 2.23, p = 0.156) and dynamics (CTb+ = 0.79 ± 0.21; CTb- = 0.55 ± 0.18; F (1,15) = 2.53, p = 0.133). Overall, these data suggest that mPFC inputs are specialized to excite reciprocally connected LH neurons.

## Discussion

Our current understanding of external inputs to the hypothalamus primarily relies on anatomical tracing studies. In addition to the lack of information about synaptic strengths, with this approach, the choice of retrograde tracer is known to significantly impact the results ^32^. These methodological constraints underscore the importance of confirming synaptic connectivity with optogenetics.

Our retrograde tracing experiments revealed significant cortical inputs to the LH from mPFC, IC, ECT, dSub and vSub. These findings are consistent with previous CTb studies in rat ^34,56^ and mouse ^57^, and reveal a marked spatial continuity of labelling across cortical areas suggesting that these projection neurons may form a more cohesive or interconnected network than previously appreciated. In addition, our findings suggest that mPFC inputs arise from a pool of neurons comparable in density to well-known subcortical inputs from the LS.

Anatomical connections from mPFC, IC and vSub have been verified previously across unrelated electrophysiological studies ^20,22,29,38^. In this study we confirmed one new pathway from ECT and showed that the anatomically labelled pathway from dSub is functionally extremely weak and likely formed mostly of fibres of passage. Importantly, we were able to compare the synaptic strength, axonal distribution, innervation density and synaptic dynamics among these pathways. When compared to inputs from the mPFC the inputs from the IC, particularly aIC, had more facilitating dynamics. This suggests that while mPFC inputs are initially powerful, IC inputs require higher frequencies. In addition, we showed that while mPFC and IC inputs are summed supralinearly in a subpopulation of neurons, neurons are differentially sensitive to each cortical channel. Input sensitivity correlated with other cellular properties suggesting the following: IC-preferring cells have elevated excitability, and strongly mPFC-recipient cells tend to project back to the mPFC.

### Local integration of monosynaptic excitatory input from the mPFC and the IC

We observed monosynaptic EPSPs consistent with the distribution of cortico-hypothalamic projections neurons in layers 5/6, which is comprised mostly of excitatory pyramidal tract and cortico-thalamic pyramidal neurons ^58^. Cortical projections are also known to reach the LH indirectly via GABAergic input from subcortical structures such as LS ^18^, CeA^59^ and the NAc ^60^, which are thought to integrate various cortical inputs prior to relaying information to the LH ^61^. Our findings, for the first time, show that LH cells can integrate direct excitatory input from the mPFC and IC locally. These inputs sum supralinearly, meaning that, when temporally aligned, their combined effect is greater than the sum of their individual contributions. Together, these findings suggest that the fastest way the cortex can influence behaviours through the LH is by exciting LH neurons. This mechanism may account for rapid sensory responses of LH neurons ^62^. The supralinear integration also implies that synchronized mPFC and IC activity could robustly drive LH neurons. Future in vivo studies should investigate whether these inputs are synchronized during behaviour and examine how the LH integrates these with inhibitory subcortical inputs. Clarifying these interactions will shed light on the broader significance of cortical and subcortical influences on LH function.

### Novel insights into the organization of cortico-hypothalamic circuits

mPFC and IC inputs to the LH displayed distinct synaptic dynamics and innervated partially overlapping domains, suggesting functional specialization of these circuits. This specialization arises, in part, through two elements of subtype selectivity: (1) electrophysiological characteristics and (2) projection target.

In our dual channel optogenetics experiment, we observed that while mPFC input was typically dominant, IC input surpassed mPFC input in a subpopulation located in dorsolateral LH. These IC-preferring neurons had elevated excitability through a hyperpolarization-activated transient current (Ih) and post-inhibitory rebound depolarization. The Ih current may allow these cells to fire bursts in vivo, potentially enabling rhythmic firing patterns ^55^. Moreover, post-inhibitory rebound makes these neurons particularly responsive after subcortical inhibition, possibly from topographically colocalized inputs from the central amygdala ^63,64^, which could modulate their activity depending on timing.

Unlike IC axons, mPFC axons were indiscriminately distributed and formed synapses with most LH neurons. Innervated LH neurons had mixed intrinsic characteristics consistent with multiple cell types ^50^. There was also marked variance in EPSP amplitude among mPFC recipient cells. While some of this response variability may stem from technical factors, such as detection limitations in distal dendrites, EPSP amplitude is also known to be actively regulated in specific subpopulations in response to conditions like cocaine exposure ^65,66^, high fat diet ^67^, and stress ^20^. Our findings are consistent with recent work showing that mPFC inputs are non-selective for several canonical LH subtypes including MCH, Orx, GABA and vGLUT2 neurons, but can modulate specific subpopulations under behaviourally relevant conditions ^20^. It is therefore likely that specialized ensembles of target-specific mPFC→LH projection neurons are required to activate specific behaviours. This complicates interpretation of average activity recording data such as fibre photometry in the mPFC→LH population and may obscure distinct behavioural contributions in opto- or chemogenetic manipulation experiments. Specialized ensembles may be reflected in the projection target of LH neurons. Our combined retrograde and anterograde experiment revealed that strongly mPFC-recipient cells tended to project back to the mPFC, suggesting a direct feedback mechanism. Reciprocal connectivity is a common feature of PFC circuits, including its interactions with the thalamus ^68^, which serve crucial functions in various cognitive processes by amplifying and sustaining representations in the PFC ^69^. In addition, several studies underscore the importance of mPFC projecting LH neurons in learning cue-food associations and cue driven consumption ^5,70,71^. Therefore, the dynamic exchange between mPFC and LH could help regulate adaptive behaviours and maintain cognitive control during complex tasks. It remains to be determined whether cortical input strength correlates with other projection targets of LH neurons ^72,73^ and if these projection patterns reflect specialized ensembles.

### Concluding remarks

Overall, these results identify the mPFC and the IC as important, rapid mediators of LH activity. The local integration of mPFC and IC inputs and the preferential reciprocal connectivity with the mPFC support the emerging view that the LH is not merely a passive recipient of cortical signals but an active computational hub capable of integrating complex information. This adds to a growing body of evidence positioning the LH as a critical player in mediating interactions between cognitive processes and adaptive behaviours ^5,6^. Further exploration of the LH’s functional architecture will be essential for uncovering the broader neurobiological mechanisms underlying survival behaviours.

## Methods

### Animals

Experiments were performed on male and female C57Bl6J mice. The animals were either obtained from Charles River or were wild-type breeding surplus from the local breeding facility. All animals were group-housed with 3 to 4 same-sex littermates and had ad libitum access to chow and water. The mice were maintained on a 12:12 light/dark cycle (lights on at 7:00 AM) in a temperature- and humidity-controlled environment. All experimental procedures were in accordance with European and Dutch law and approved by the central committee animal experiments and local animal ethical care committee (IvD) of the VU University and VU University Medical Centre (Amsterdam, Netherlands).

### Surgeries

To achieve anaesthesia and analgesia, mice received an intraperitoneal (i.p.) injection of Fentanyl (0.05 mg/kg), Medetomidine (0.5 mg/kg), and Midazolam (5 mg/kg) in saline prior to surgery. The mice were then placed in a stereotaxic frame with adjustable ear bars. To prevent drying, the eyes were covered with ointment. Lidocaine (10 mg/kg) was administered under the scalp for additional local analgesia. After exposing the skull, small craniotomies were made above the coordinates for the injections (see Table 1 for injection coordinates). Glass injection needles were filled with the injectate (retrograde tracers and/or virus diluted in sterile PBS; see Table 1 for specific concentrations) and inserted into the brain. The injectate was delivered using a Narishige microinjector at a rate of 10 nl per minute for LH injections or 30 nl per minute for all other injections. The needle was left in place for 25 minutes after LH injections and 5 minutes for all other injections. Upon completion of the injections, the skin was closed with sutures. The animals were then removed from the stereotaxic frame and received an i.p. injection of Flumazenil (0.5 mg/kg) and Atipamezole (2.5 mg/kg) to antagonize the anaesthesia. The mice were placed back into their home cages on a heating pad and allowed to recover. Drinking water was supplemented with carprofen one day prior to surgery and continued for 3 to 5 days post-surgery.

### Histology

Tissue was collected 7-10 days after CTb injections and 3-4 weeks after anterograde tracing surgeries to allow for viral expression. Mice were euthanized with a terminal dose of pentobarbital and perfused transcardially with PBS followed by 4% paraformaldehyde (PFA). Brains were extracted and post-fixed in PFA for at least 48 hours. Brains were then sliced at 50-70 µm thickness and mounted using Mowiol mounting medium containing 5 µM DAPI for nuclear counterstaining. Slices were imaged with a Vectra Polaris slide scanner (Akoya Biosciences) at 20x resolution.

### Retrograde tracing

Retrogradely labelled cells were quantified semi-automatically using QuPath software. Regions of interest were manually annotated using fiducial landmarks and the Allen Brain Atlas. Cell counting was performed using a combination of manual detection and automatic detection via QuPath’s built-in cell detection function.

For LH retrograde tracing (experiment 1), three brains were analysed from AP coordinates 2.23 mm to -3.79 from bregma. Additionally, to generate a 3D model of long-range projections to the LH, one brain was registered to the Allen Brain Atlas using the ABBA software package ^74^. Detected cells were mapped onto a 3D brain model using Brainrender ^75^.

In combined retrograde and anterograde tracing experiments to characterize reciprocal connections (experiment 4) between the IC-LH (n=2 brains) and mPFC-LH (n=2 brains), retrogradely labelled cells in the LH were quantified in brain sections spanning -1.07 mm to 2.06 mm from bregma.

### Anterograde tracing

To quantify the localization and density of axonal fibres in the hypothalamus, AAV1-hSyn-ChR2-YFP was injected into either mPFC or IC (see Table 1 for coordinates). Brain tissue was processed as previously described. Coronal sections containing the LH at anteroposterior coordinates of -1.07, -1.31, -1.55, and -1.79 mm from bregma were analysed for three brains.

For comparative analysis of fibre distribution, images were thresholded and binarized. A custom ImageJ macro was used to overlay a 20x20 grid on the LH, with grid alignment based on fiducial landmarks (fornix, top of the ventricle, bottom of the section, and lateral edge of the LH). For each section, average intensity values within the grid were measured in ImageJ and exported as arrays to MATLAB. In MATLAB, kernel density estimation was performed to determine the distribution of thresholded pixels across the mediolateral and dorsoventral axes. Intensity measurements and density distributions were averaged for each anteroposterior location and injection group.

### Ex vivo patch-clamp electrophysiology

#### Brain slice preparation and solutions

3-4 weeks after viral injections, mice were euthanized by cervical dislocation under isoflurane anaesthesia. Brains were quickly removed following decapitation and placed in ice-cold N-methyl-D-glucamine (NMDG) solution, which included (in mM): 2.5 KCl, 1.2 NaH2PO4, 30 NaHCO3, 20 HEPES, 25 Glucose, 5 Na ascorbate, 3 Na pyruvate, 93 NMDG, 10 MgSO4, and 0.5 CaCl2 (pH 7.3-7.4, 300-310 mOsm). Coronal slices (250 µm thick) containing the lateral hypothalamus (LH) and injection location were prepared using a vibratome (Leica VS1000). The slices were collected and placed in a holding chamber with the NMDG solution maintained at 34°C. After 20 minutes, the slices were transferred to another holding chamber containing artificial cerebrospinal fluid (ACSF) and allowed to recover for at least 1 hour at room temperature before recordings. The ACSF solution included (in mM): 126 NaCl, 3 KCl, 2 MgSO4, 1.1 NaH2PO4, 2 CaCl2, 10 Glucose, 0.1 Na-pyruvate, 0.4 ascorbic acid, 0.5 glutamine, and 26 NaHCO3 (pH 7.3-7.4 and 300-315 mOsm). During slicing, recovery, and recordings, the solutions were continuously bubbled with 95% O2 and 5% CO2. To verify viral targeting, slices containing the injection location were post-fixed in 4% PFA for 3-5 days, then mounted with Mowiol mounting medium with 5 µM DAPI for nuclear counterstain. Mice were only included in the analysis if EYFP expression was restricted to the injection location.

#### Recording and analysis

For electrophysiological recordings, brain slices were placed in a recording chamber that was continuously perfused with oxygenated ACSF maintained at 32°C. Neurons in the LH were visualized using an Olympus BX51WI upright microscope equipped with infrared differential interference contrast optics. Patch pipettes, with a resistance of 3-6 MΩ, were pulled from borosilicate glass capillaries and filled with an internal solution based on potassium-gluconate, containing in (in mM): 130 K-gluconate, 5 NaCl, 2 MgSO4, 10 HEPES, 0.1 EGTA, 4 MgATP, 0.4 Na-GTP, and 2 Pyruvic Acid (pH adjusted to 7.3 with KOH). Recordings were performed using Multiclamp 700B amplifiers (Axon Instruments, Molecular Devices). The data were filtered at 3 kHz, digitized at 10 kHz (using a National Instruments USB-6343 digitizer), and collected with MIES software (https://github.com/AllenInstitute/MIES) running in IgorPro 8 (WaveMetrics). Recordings were included if access resistance (monitored with a 5mV hyperpolarizing step between stimulation sweeps) was <25 MΩ and did not increase more than 20%, and if the net leak current did not exceed -500pA.

#### Optogenetics

Photostimulation of expressed opsin(s) was achieved by delivering 470 nm and or 590 nm light through a 40x objective using an LED system (DC4100; Thorlabs Inc, USA).

### Channelrhodopsin-2

In experiments using ChR2, blue irradiance was set at 0.7-1 mW/mm^2^. Cells were current-clamped around -65 mV. To assess action potentials occurring within these stimulus trains, sweeps with action potentials prior to light stimulation were excluded. A train of 5 blue light pulses of 3 ms each was delivered at 10 Hz and repeated every 15 seconds to evoke optogenetic responses. At least 7 sweeps were averaged to calculate EPSP parameters, excluding those with action potentials. The response threshold was set at 4 * the standard deviation of a baseline, computed prior to each pulse. Responses exceeding the response threshold were considered optically evoked responses and included for further analysis. Latency was defined as time from the light pulse onset to passing the response threshold. The progressive paired pulse ratio was computed by dividing the amplitude to each pulse by the amplitude to the first light pulse of stimulus trains.

### ChrimsonR and Chronos

In experiments using ChrimsonR and or Chronos (experiment 3), delivery of 470 nm and 590 nm light was achieved using a mirror without additional filters. In validation experiments, neurons were recorded at the injection site of animals injected only with ChrimsonR or Chronos. Opsin expressing neurons were current-clamped and stimulated with 10 Hz trains of 5 pulses of 3 ms to get a firing probability across a range of increasing irradiances of 470 and 590 nm light. Results were similar to recordings from brains with both opsins present, where any possible postsynaptic effects were eliminated by DNQX – these data were therefore combined.

In animals only expressing ChrimsonR, LH cells were voltage clamped at -70 mV to assess postsynaptic responses across a range of different light irradiances for 590 nm and 470 nm light. A sequential photostimulation protocol was used (Fig. 3 and Fig. S5) to suppress observed crosstalk of ChrimsonR with 470 nm photostimulation. Photostimulation protocols were as follows: 1) 10 ms 590 nm, 2) 10 ms 470 nm, 3) 250 ms 590 nm followed by 10 ms 470 nm light.

In subsequent experiments both ChrimsonR and Chronos were expressed in the mPFC and IC of each animal (mPFC_ChrimsonR_ - IC_Chronos_ or mPFC_Chronos_ – IC_ChrimsonR_). The sequential photostimulation protocols described above were used to categorize responses. To compare amplitudes, the 590 nm 10 ms light pulse was always compared to the desensitized 470 nm 10 ms light pulse. To study integration of inputs, these individual responses were summed and compared to the amplitude of simultaneous stimulation (10 ms 470 nm and 590 nm).

Input resistance, voltage sag and membrane tau were calculated from hyperpolarizing steps. PIR was defined as a depolarization of >5mV compared to baseline at the end of these hyperpolarizing steps. In addition, a current ramp (0-200 pA; duration 15 s) was injected to study potential differences in firing rate. Firing rate was calculated within 8 pA bins (200 bins, 600 ms per bin). Rheobase was defined as the minimal current required to elicit the first AP. The gain was defined for each neuron as the slope of the linear fit between the bin containing the rheobase and the bin containing the max firing frequency.

Amplitudes and frequency of spontaneous events (sEPSCs) occurring between photostimulation were measured. For each cell the empirical cumulative distribution function (ecdf) was calculated for the amplitudes of spontaneous events, were a value x in the function F(x) corresponds to the proportion of the values less than or equal to x. P(oEPSC ≥ sEPSCs) was defined as the approximation of the ecdf for the oEPSC amplitude, and indicates the proportion of sEPCS amplitudes equal to or smaller than the oEPSC amplitude. In other words, a P (oEPSC ≥ sEPSCs) of 70% indicates that the observed oEPSC amplitude is larger than 70% of the spontaneous events.

#### Pharmacology

In certain experiments, the voltage-gated sodium channel blocker Tetrodoxin (TTX; 1 µM) was added to the ACSF to inhibit action potentials, along with the voltage-gated potassium channel blocker 4-Aminopyridine (4-AP; 250 µM). As described above, DNQX (10 µM) was added to the ACSF during recordings of opsin expressing neurons in the mPFC and IC, to block excitatory connections between these areas in cases when both opsins were injected into the same animal. All pharmacological agents were bath-applied for at least 5 minutes before recordings.

#### Locations of recorded neurons

To capture locations of recorded neurons an image of pipette placement within the lateral hypothalamus was captured after each recording with a 4x objective. Locations relative to the boundaries of the lateral hypothalamus were then calculated using Fiji software.

### Data analysis & statistics

Data analysis and visualization were performed using custom scripts in ImageJ, MATLAB (2019b), Python, and QuPath. All statistical analyses were conducted in MATLAB (2019b). Detailed descriptions of the specific statistical tests are provided in Table S2.

Normality was assessed using the Shapiro-Wilk test before statistical testing. For data that did not meet normality assumptions, non-parametric tests were applied, including the Wilcoxon signed-rank test, Wilcoxon rank-sum test, and Kruskal-Wallis test. Multiple comparisons were corrected using the false discovery rate (FDR) method. Proportional data, such as response rates, were analysed using Chi-square tests and followed up with Fisher’s Exact tests.

Generalized linear mixed effects models (GLMEs) were applied to non-normally distributed multifactorial datasets and analyses involving repeated measures (e.g., accounting for opsin and cortical area effects on inputs in dual-channel experiments or comparing responses to consecutive pulses). Model selection was based on visual inspection of data distributions to identify the appropriate distribution family (e.g., Gamma), along with model performance evaluation using the Akaike Information Criterion (AIC) and residual analysis.

All data are presented as mean ± SEM unless otherwise specified. Statistical significance is indicated in figures using the following notation: p < 0.05 (*), p < 0.01 (**), or p < 0.001 (***).

## Author contributions

**L.J.A.M. Razenberg**: Conceptualization, Methodology, Formal analysis, Investigation, Visualization, Supervision, Writing – Original draft. **P. de Greef**: Data Curation, Investigation. **H.D. Mansvelder**: Funding acquisition, Supervision, Writing – Review & Editing. **M.M. Karnani**: Conceptualization, Methodology, Supervision, Writing - Review & Editing.

## Acknowledgements

We thank Tim Heistek and Jaap Timmermans for excellent technical assistance and the Microscopy Unit from the Molecular Cell Biology & Immunology Technology Centre, in particular Marko Popovic, for use and maintenance of the slide-scanner. We also thank Joke Wortel and Rolinka van der Loo for their support with animal care.

