## Supplementary Table S2 for "Integration of cortical inputs in the lateral hypothalamus is dominated by the medial prefrontal cortex"

| Figure | Variable | Groups | n | Mean | SEM | Test | | | | | p | | Pairwise comparisons | | | Statistic | | Adjusted-p |
|---|---|---|---|---|---|---|---|---|---|---|---|---|---|---|---|---|---|---|
| Fig 2 | | | | | | | | | | | | | | | | | | |
| 2D | Response rate | mPFC | 49/79 | 62.03% | | Chi Square | | df | n | Chi-sq | p | | Fisher's Exact tests | | | OddsRatio | p | fdr |
| | | aIC | 10/29 | 34.48% | | | | 6 | 265 | 46.44 | | **0.000** | | mPFC | aIC | 3.10 | 0.016 | **0.025** |
| | | -IC- | 20/68 | 29.41% | | | | | | | | | | mPFC | -IC- | 3.92 | 0.000 | **0.000** |
| | | pIC | 6/25 | 24.00% | | | | | | | | | | mPFC | pIC | 5.17 | 0.001 | **0.003** |
| | | ECT | 2/20 | 10.00% | | | | | | | | | | mPFC | ECT | 14.70 | 0.000 | **0.000** |
| | | dSub | 1/29 | 3.45% | | | | | | | | | | mPFC | dSub | 45.73 | 0.000 | **0.000** |
| | | vSub | 4/15 | 26.67% | | | | | | | | | | mPFC | vSub | 4.49 | 0.021 | **0.028** |
| | | | | | | | | | | | | | | aIC | -IC- | 1.26 | 0.638 | 0.408 |
| | | | | | | | | | | | | | | aIC | pIC | 1.67 | 0.552 | 0.400 |
| | | | | | | | | | | | | | | aIC | ECT | 4.74 | 0.089 | 0.088 |
| | | | | | | | | | | | | | | aIC | dSub | 14.74 | 0.005 | **0.010** |
| | | | | | | | | | | | | | | aIC | vSub | 1.45 | 0.738 | 0.446 |
| | | | | | | | | | | | | | | -IC- | pIC | 1.32 | 0.795 | 0.455 |
| | | | | | | | | | | | | | | -IC- | ECT | 3.75 | 0.139 | 0.126 |
| | | | | | | | | | | | | | | -IC- | dSub | 11.67 | 0.003 | **0.007** |
| | | | | | | | | | | | | | | -IC- | vSub | 1.15 | 1.000 | 0.518 |
| | | | | | | | | | | | | | | pIC | ECT | 2.84 | 0.269 | 0.225 |
| | | | | | | | | | | | | | | pIC | dSub | 8.84 | 0.041 | **0.044** |
| | | | | | | | | | | | | | | pIC | vSub | 0.87 | 1.000 | 0.544 |
| | | | | | | | | | | | | | | ECT | dSub | 3.11 | 0.559 | 0.380 |
| | | | | | | | | | | | | | | ECT | vSub | 0.31 | 0.367 | 0.285 |
| | | | | | | | | | | | | | | dSub | vSub | 0.10 | 0.039 | **0.047** |
| 2E | Amplitude P1 | mPFC | 49 | 3.73 ± | 0.55 | Kruskal Wallis | | | | | | | Wilcoxon rank sum | | | | | |
| | | aIC | 10 | 0.81 ± | 0.16 | 'Source' | 'SS' | 'df' | 'MS' | 'Chi-sq' | p | | | mPFC | aIC | 1362.00 | 0.001 | **0.016** |
| | | -IC- | 20 | 1.89 ± | 0.79 | 'Groups' | 11703.96 | 6 | 1950.66 | 18.77 | | **0.005** | | mPFC | -IC- | 1611.00 | 0.002 | **0.025** |
| | | pIC | 6 | 2.15 ± | 1.04 | 'Error' | 41293.54 | 79 | 522.703 | | | | | mPFC | pIC | 1168.00 | 0.174 | 0.545 |
| | | ECT | 2 | 3.08 ± | 1.09 | 'Total' | 52997.50 | 85 | | | | | | mPFC | ECT | 1029.00 | 0.808 | 0.849 |
| | | dSub | 1 | 5.11 | | | | | | | | | | mPFC | dSub | 1000.00 | 0.376 | 0.824 |
| | | vSub | 4 | 3.10 ± | 2.15 | | | | | | | | | mPFC | vSub | 1099.00 | 0.444 | 0.824 |
| | | | | | | | | | | | | | | aIC | -IC- | 139.00 | 0.630 | 0.824 |
| | | | | | | | | | | | | | | aIC | pIC | 80.00 | 0.635 | 0.824 |
| | | | | | | | | | | | | | | aIC | ECT | 55.00 | 0.030 | 0.212 |
| | | | | | | | | | | | | | | aIC | dSub | 55.00 | 0.182 | 0.545 |
| | | | | | | | | | | | | | | aIC | vSub | 68.00 | 0.374 | 0.824 |
| | | | | | | | | | | | | | | -IC- | pIC | 239.00 | 0.633 | 0.824 |
| | | | | | | | | | | | | | | -IC- | ECT | 196.00 | 0.134 | 0.545 |
| | | | | | | | | | | | | | | -IC- | dSub | 191.00 | 0.165 | 0.545 |
| | | | | | | | | | | | | | | -IC- | vSub | 223.00 | 0.715 | 0.834 |
| | | | | | | | | | | | | | | pIC | ECT | 24.00 | 0.429 | 0.824 |
| | | | | | | | | | | | | | | pIC | dSub | 22.00 | 0.571 | 0.824 |
| | | | | | | | | | | | | | | pIC | vSub | 32.00 | 0.914 | 0.914 |
| | | | | | | | | | | | | | | ECT | dSub | 3.00 | 0.667 | 0.824 |
| | | | | | | | | | | | | | | ECT | vSub | 9.00 | 0.533 | 0.824 |
| | | | | | | | | | | | | | | dSub | vSub | 4.00 | 0.800 | 0.849 |
| - | Max Amplitude | mPFC | 49 | 4.28 ± | 0.19 | Kruskal Wallis | | | | | | | | | | | | |
| | | aIC | 10 | 1.68 ± | 0.43 | 'Source' | 'SS' | 'df' | 'MS' | 'Chi-sq' | p | | | | | | | |
| | | -IC- | 20 | 2.71 ± | 0.34 | 'Groups' | 6571.06 | 6 | 1095.176 | 10.54 | | 0.104 | | | | | | |
| | | pIC | 6 | 2.36 ± | 0.16 | 'Error' | 46426.44 | 79 | 587.6765 | | | | | | | | | |
| | | ECT | 2 | 4.51 ± | 0.34 | 'Total' | 52997.50 | 85 | | | | | | | | | | |
| | | dSub | 1 | 5.11 | | | | | | | | | | | | | | |
| | | vSub | 4 | 4.50 ± | 0.58 | | | | | | | | | | | | | |
| 2F | Latency | mPFC | 49 | 5.64 ± | 0.19 | Kruskal Wallis | | | | | | | | | | | | |
| | | aIC | 10 | 6.51 ± | 0.43 | 'Source' | 'SS' | 'df' | 'MS' | 'Chi-sq' | p | | | | | | | |
| | | -IC- | 20 | 6.46 ± | 0.34 | 'Groups' | 6987.59 | 6 | 1164.599 | 11.47 | | 0.075 | | | | | | |

| | | | | | | | | | | | | | | | | | |
|---|---|---|---|---|---|---|---|---|---|---|---|---|---|---|---|---|---|
| | | pIC | 6 | 6.67 ± | 0.16 | 'Error' | 44182.41 | 78 | 566.4411 | | | | | | | | |
| | | ECT | 2 | 5.22 ± | 0.34 | 'Total' | 51170.00 | 84 | | | | | | | | | |
| | | dSub | 1 | 5.85 | | | | | | | | | | | | | |
| | | vSub | 4 | 4.93 ± | 0.58 | | | | | | | | | | | | |
| 2G | PPR | mPFC | | | | GLME | PPR ~ Group * Pulsenr + (1\|cellid)' | | | | | | | | | | |
| | | aIC | | | | Term | Fstat | DF1 | DF2 | | p | | | | | | |
| | | -IC- | | | | Intercept | 62.72 | 1 | 395 | | | 0.000 | | | | | |
| | | pIC | | | | Group | 0.22 | 6 | 396 | | | 0.970 | | | | | |
| | | ECT | | | | PulseNr | 3.95 | 4 | 396 | | | **0.004** | | | | | |
| | | dSub | | | | Group * PulseNr | 1.65 | 24 | 396 | | | **0.030** | | | | | |
| | | vSub | | | | | | | | | | | | | | | |
| 2H | P5/P1 | mPFC | 49 | 0.89 ± | 0.15 | Kruskal Wallis | | | | | | | Wilcoxon rank sum | | ranksum | p | fdr |
| | | aIC | 10 | 2.12 ± | 0.55 | 'Source' | 'SS' | 'df' | 'MS' | 'Chi-sq' | p | | mPFC | aIC | 1063.00 | 0.001 | **0.014** |
| | | -IC- | 20 | 2.00 ± | 0.50 | 'Groups' | 12851.45 | 6 | 2141.909 | 20.61 | | **0.002** | mPFC | -IC- | 1207.00 | 0.003 | **0.017** |
| | | pIC | 6 | 0.97 ± | 0.38 | 'Error' | 40146.05 | 79 | 508.1778 | | | | mPFC | pIC | 1111.00 | 0.754 | 0.558 |
| | | ECT | 2 | 0.85 ± | 0.67 | 'Total' | 52997.50 | 85 | | | | | mPFC | ECT | 1036.00 | 0.936 | 0.654 |
| | | dSub | 1 | 0.28 ± | 0.00 | | | | | | | | mPFC | dSub | 1027.00 | 0.264 | 0.369 |
| | | vSub | 4 | 1.57 ± | 0.23 | | | | | | | | mPFC | vSub | 1018.00 | 0.026 | 0.111 |
| | | | | | | | | | | | | | aIC | -IC- | 170.00 | 0.371 | 0.389 |
| | | | | | | | | | | | | | aIC | pIC | 103.00 | 0.056 | 0.176 |
| | | | | | | | | | | | | | aIC | ECT | 70.00 | 0.364 | 0.416 |
| | | | | | | | | | | | | | aIC | dSub | 65.00 | 0.182 | 0.286 |
| | | | | | | | | | | | | | aIC | vSub | 76.00 | 0.945 | 0.626 |
| | | | | | | | | | | | | | -IC- | pIC | 270.00 | 0.152 | 0.383 |
| | | | | | | | | | | | | | -IC- | ECT | 216.00 | 0.436 | 0.392 |
| | | | | | | | | | | | | | -IC- | dSub | 208.00 | 0.165 | 0.347 |
| | | | | | | | | | | | | | -IC- | vSub | 219.00 | 0.491 | 0.411 |
| | | | | | | | | | | | | | pIC | ECT | 27.00 | 1.000 | 0.629 |
| | | | | | | | | | | | | | pIC | dSub | 26.00 | 0.571 | 0.449 |
| | | | | | | | | | | | | | pIC | vSub | 26.00 | 0.171 | 0.308 |
| | | | | | | | | | | | | | ECT | dSub | 4.00 | 1.000 | 0.599 |
| | | | | | | | | | | | | | ECT | vSub | 4.00 | 0.267 | 0.335 |
| | | | | | | | | | | | | | dSub | vSub | 1.00 | 0.400 | 0.387 |
| 2I | Facilitation rate | mPFC | 11/49 | 22.45% | | Chi Square | | | | | | | Fisher's Exact tests | | OddsRatio | p | fdr |
| | | aIC | 7/10 | 70.00% | | | | df | n | Chi-sq | p | | mPFC | aIC | 0.12 | 0.006 | 0.117 |
| | | -IC- | 11/20 | 55.00% | | | | 6 | 92 | 14.94 | | **0.021** | mPFC | -IC- | 0.24 | 0.012 | 0.115 |
| | | pIC | 2/6 | 33.33% | | | | | | | | | mPFC | pIC | 0.58 | 0.619 | 1.000 |
| | | ECT | 1/2 | 50.00% | | | | | | | | | mPFC | ECT | 0.29 | 0.419 | 1.000 |
| | | dSub | 0/1 | 0.00% | | | | | | | | | mPFC | dSub | 65535.00 | 1.000 | 1.000 |
| | | vSub | 3/4 | 75.00% | | | | | | | | | mPFC | vSub | 0.10 | 0.052 | 0.337 |
| | | | | | | | | | | | | | aIC | -IC- | 1.91 | 0.694 | 1.000 |
| | | | | | | | | | | | | | aIC | pIC | 4.67 | 0.302 | 1.000 |
| | | | | | | | | | | | | | aIC | ECT | 2.33 | 1.000 | 1.000 |
| | | | | | | | | | | | | | aIC | dSub | 65535.00 | 0.364 | 1.000 |
| | | | | | | | | | | | | | aIC | vSub | 0.78 | 1.000 | 1.000 |
| | | | | | | | | | | | | | -IC- | pIC | 2.44 | 0.645 | 1.000 |
| | | | | | | | | | | | | | -IC- | ECT | 1.22 | 1.000 | 1.000 |
| | | | | | | | | | | | | | -IC- | dSub | 65535.00 | 0.476 | 1.000 |
| | | | | | | | | | | | | | -IC- | vSub | 0.41 | 0.615 | 1.000 |
| | | | | | | | | | | | | | pIC | ECT | 0.50 | 1.000 | 1.000 |
| | | | | | | | | | | | | | pIC | dSub | 65535.00 | 1.000 | 0.975 |
| | | | | | | | | | | | | | pIC | vSub | 0.17 | 0.524 | 1.000 |
| | | | | | | | | | | | | | ECT | dSub | 65535.00 | 1.000 | 0.929 |
| | | | | | | | | | | | | | ECT | vSub | 0.33 | 1.000 | 1.000 |
| | | | | | | | | | | | | | dSub | vSub | 0.00 | 0.400 | 1.000 |

| | | | | | | | | | | | | | | | | | | |
|---|---|---|---|---|---|---|---|---|---|---|---|---|---|---|---|---|---|---|
| 2K | AP probability (per pulse) | mPFC | | | | | GLME | prob ~ group * Pulsenr + (1\|cellID) | | | | | | | | | | |
| | | aIC | | | | | Term | Fstat | Df1 | Df2 | | p | | | | | | |
| | | -IC- | | | | | Intercept | 470.67 | 1 | 355 | | **0.000** | | | | | | |
| | | pIC | | | | | group | 0.82 | 6 | 355 | | 0.552 | | | | | | |
| | | ECT | | | | | Pulsenr | 1.99 | 4 | 355 | | 0.096 | | | | | | |
| | | dSub | | | | | group:Pulsenr | 0.82 | 24 | 355 | | 0.714 | | | | | | |
| | | vSub | | | | | | | | | | | | | | | | |
| 2L | AP probability (1 or more AP) | mPFC | 49 | 11.75% | ± | 0.04 | Kruskal Wallis | | | | | | | | | | | |
| | | aIC | 10 | 0.00% | ± | 0.00 | 'Source' | 'SS' | 'df' | 'MS' | 'Chi-sq' | p | | | | | | |
| | | -IC- | 20 | 1.85% | ± | 0.02 | 'Groups' | 960.27 | 6 | 160.0443 | 5.54 | 0.476 | | | | | | |
| | | pIC | 6 | 6.67% | ± | 0.06 | 'Error' | 12376.23 | 71 | 174.3132 | | | | | | | | |
| | | ECT | 2 | 0.00% | ± | 0.00 | 'Total' | 13336.50 | 77 | | | | | | | | | |
| | | dSub | 1 | 0.00% | ± | 0.00 | | | | | | | | | | | | |
| | | vSub | 4 | 0.00% | ± | 0.00 | | | | | | | | | | | | |
| Fig 3 | | | | | | | | | | | | | | | | | | |
| 3D | Desensitization | ChrimsonR$_{LED590}$ | 19 | -25.99 | ± | 4.99 | | | | | | | **Wilcoxon rank sum test** | | | Z | p | FDR |
| | | ChrimsonR$_{LED470}$ | 19 | -18.50 | ± | 3.72 | | | | | | | ChrimsonR | Chronos | LED590 | -2.9052 | 0.004 | **0.011** |
| | | ChrimsonR$_{LED590-D}$ | 19 | 0.00 | ± | 0.00 | | | | | | | Chrimson | Both | LED590 | -1.5295 | 0.126 | 0.227 |
| | | Chronos$_{LED590}$ | 6 | 0.00 | ± | 0.00 | | | | | | | Chronos | Both | LED590 | 2.35 | 0.019 | **0.042** |
| | | Chonos$_{LED470}$ | 6 | -17.30 | ± | 4.10 | | | | | | | ChrimsonR | Chronos | LED470 | 0.28 | 0.777 | 0.777 |
| | | Chronos$_{LED470-D}$ | 6 | -10.74 | ± | 2.75 | | | | | | | Chrimson | Both | LED471 | 0.75 | 0.452 | 0.678 |
| | | Both$_{LED590}$ | 15 | -40.32 | ± | 20.22 | | | | | | | Chronos | Both | LED472 | 0.48 | 0.633 | 0.777 |
| | | Both$_{LED470}$ | 15 | -36.31 | ± | 13.22 | | | | | | | ChrimsonR | Chronos | Desentizit | 3.88 | 0.000 | **0.000** |
| | | Both$_{LED470-D}$ | 15 | -12.09 | ± | 2.69 | | | | | | | Chrimson | Both | Desentizit | 4.66 | 0.000 | **0.000** |
| | | | | | | | | | | | | | Chronos | Both | Desentizit | -0.3717 | 0.710 | 0.777 |
| | *EPSC parameters* | | | | | | GLME | *var* ~ opsin*cortex + (1\| cell) | | | | | | | | | | |
| 3E | Amplitude (pA) | mPFC$_{ChrimsonR}$ | 29 | 32.08 | ± | 9.96 | Term | Fstat | DF1 | DF2 | | p | | | | | | |
| | | mPFC$_{Chronos}$ | 29 | 12.40 | ± | 4.28 | Intercept | 6.0559 | 1 | 86.00 | | **0.016** | | | | | | |
| | | IC$_{ChrimsonR}$ | 16 | 4.11 | ± | 1.51 | Cortex | 4.0175 | 1 | 86.00 | | **0.048** | | | | | | |
| | | IC$_{Chronos}$ | 16 | 5.72 | ± | 1.69 | Opsin | 0.4521 | 1 | 86.00 | | 0.503 | | | | | | |
| | | | | | | | Cortex:Opsin | 0.032 | 1 | 86.00 | | 0.858 | | | | | | |
| 3F | Latency (ms) | mPFC$_{ChrimsonR}$ | 29 | 3.91 | ± | 0.32 | Intercept | 245.67 | 1 | 51.00 | | **0.000** | | | | | | |
| | | mPFC$_{Chronos}$ | 29 | 3.85 | ± | 0.53 | Cortex | 7.1289 | 1 | 51.00 | | **0.010** | | | | | | |
| | | IC$_{ChrimsonR}$ | 16 | 5.31 | ± | 0.67 | Opsin | 0.10993 | 1 | 51.00 | | 0.742 | | | | | | |
| | | IC$_{Chronos}$ | 16 | 6.04 | ± | 0.59 | Cortex:Opsin | 0.011865 | 1 | 51.00 | | 0.914 | | | | | | |
| 3G | Rise time (ms) | mPFC$_{ChrimsonR}$ | 29 | 2.52 | ± | 0.29 | Intercept | 21.469 | 1 | 49.00 | | **0.000** | | | | | | |
| | | mPFC$_{Chronos}$ | 29 | 1.83 | ± | 0.33 | Cortex | 5.0707 | 1 | 49.00 | | **0.029** | | | | | | |
| | | IC$_{ChrimsonR}$ | 16 | 3.57 | ± | 0.69 | Opsin | 0.66019 | 1 | 49.00 | | 0.420 | | | | | | |
| | | IC$_{Chronos}$ | 16 | 4.56 | ± | 0.76 | Cortex:Opsin | 0.15678 | 1 | 49.00 | | 0.694 | | | | | | |
| 3H | Decay time (ms) | mPFC$_{ChrimsonR}$ | 29 | 9.92 | ± | 1.47 | Intercept | 118.73 | 1 | 49.00 | | **0.000** | | | | | | |
| | | mPFC$_{Chronos}$ | 29 | 4.23 | ± | 0.86 | Cortex | 0.017778 | 1 | 49.00 | | 0.894 | | | | | | |
| | | IC$_{ChrimsonR}$ | 16 | 5.93 | ± | 0.82 | Opsin | 2.0335 | 1 | 49.00 | | 0.160 | | | | | | |
| | | IC$_{Chronos}$ | 16 | 10.65 | ± | 1.87 | Cortex:Opsin | 1.013 | 1 | 49.00 | | 0.319 | | | | | | |
| | *Response integration* | | | | | | Wilcoxon signed rank test | | | | | | | | | | | |
| 3K | Amplitude (pA) | Summed | 10 | 66.17 | ± | 27.19 | Normalized to summed response | | n | zval | | p | | | | | | |

| | | | | | | | | | | | |
|---|---|---|---|---|---|---|---|---|---|---|---|
| | | Dual | 10 | 91.74 ± | 45.65 | Summed vs. Dual Stim | | 10 | -2.2934 | | **0.022** |
| 3L | Surface area (pA * s) | Summed | 10 | 0.55 ± | 0.21 | Normalized to summed response | | 10 | -2.0896 | | **0.037** |
| | | Dual | 10 | 0.74 ± | 0.32 | Summed vs. Dual Stim | | | | | |
| Fig 4 | | | | | | | | | | | |
| 4A | mPFC EPSC Amplitude | All | 40 | 25.08 ± | 6.70 | | | | | | |
| | | $mPFC_{Preferring}$ | 30 | 36.35 ± | 9.43 | | | | | | |
| | | $IC_{Preferring}$ | 10 | 3.81 ± | 1.23 | | | | | | |
| | IC EPSC Amplitude | All | 40 | 4.68 ± | 1.14 | | | | | | |
| | | $mPFC_{Preferring}$ | 30 | 2.2328 ± | 0.79 | | | | | | |
| | | $IC_{Preferring}$ | 10 | 14.36 ± | 2.99 | | | | | | |
| | | | | | | Wilcoxon Signed rank tests | | | | | |
| 4B | EPSC Amplitude | | | | | Normalized to max input | | n | zval | p | |
| | $mPFC_{Preferring}$ | mPFC EPSC | 30 | 1.00 ± | 0.00 | mPFC vs. IC EPSC | | 30 | 4.99 | | **0.000** |
| | | IC EPSC | 30 | 0.11 ± | 0.04 | | | | | | |
| 4D | EPSC Amplitude | | | | | Normalized to max input | | n | zval | p | |
| | $IC_{Preferring}$ | mPFC EPSC | 10 | 0.31 ± | 0.11 | mPFC vs. IC EPSC | | 10 | -2.8214 | | **0.005** |
| | | IC EPSC | 10 | 1.00 ± | 0.00 | | | | | | |
| 4G | Rinput | $mPFC_{Preferring}$ | 21 | 344.99 ± | 7.77 | Wilcoxon rank sum | n1 | n2 | zval | p | |
| | | $IC_{Preferring}$ | 9 | 371.09 ± | 11.88 | | 21 | 9 | -0.4526 | | 0.651 |
| 4H | Sag | $mPFC_{Preferring}$ | 20 | 4.29 ± | 0.24 | Wilcoxon rank sum | n1 | n2 | zval | p | |
| | | $IC_{Preferring}$ | 9 | 10.30 ± | 0.93 | | 20 | 9 | -2.2392 | | **0.025** |
| 4I | PIR | $mPFC_{Preferring}$ | 9/20 | 45% | | Chi Square test | df | n | Chi-sq | p | |
| | | $IC_{Preferring}$ | 7/8 | 88% | | | 1 | 28 | 5.32 | | **0.021** |
| 4J | *Cell location* | | | | | Wilcoxon rank sum | n1 | n2 | zval | p | |
| | Medial-Lateral | $mPFC_{Preferring}$ | 30 | 0.44 ± | 0.08 | | 30 | 10 | -0.7965 | | 0.426 |
| | | $IC_{Preferring}$ | 10 | 0.61 ± | 0.14 | | | | | | |
| | Dorsal-Ventral | $mPFC_{Preferring}$ | 30 | 0.43 ± | 0.04 | | 30 | 10 | -1.9834 | | **0.047** |
| | | $IC_{Preferring}$ | 10 | 0.57 ± | 0.06 | | | | | | |
| | Anterior-Posterior | $mPFC_{Preferring}$ | 30 | -1.39 ± | 0.05 | | 30 | 10 | 0.48 | | 0.634 |
| | | $IC_{Preferring}$ | 10 | -1.42 ± | 0.08 | | | | | | |
| Fig 5 | | | | | | | | | | | |
| 5C | CTb counts | mPFC(total) | 2 | 58.50 ± | 1.50 | GLME | counts ~ Group * AP + (1 \| sample)' | | | | |
| | | IC(total) | 2 | 15.00 ± | 6.00 | Term | Fstat | DF1 | DF2 | p | |
| | | | | | | Intercept | 12.1414 | 1 | 32.00 | | **0.002** |
| | | | | | | Group | 11.2795 | 1 | 32.00 | | **0.002** |
| | | | | | | AP | 2.0197 | 1 | 32.00 | | 0.165 |
| | | | | | | Group*AP | 5.8567 | 1 | 32.00 | | **0.021** |
| 5G | Response rate | $mPFC_{CTB+}$ | 11/11 | 100% | | Chi Square test | df | n | Chi-sq | p | |
| | | $mPFC_{CTB-}$ | 10/15 | 67% | | mPFC | 1 | 26 | 4.54 | | **0.033** |
| | | $IC_{CTB+}$ | 2/6 | 33% | | IC | 1 | 16 | 0.02 | | 0.889 |
| | | $IC_{CTB-}$ | 3/10 | 30% | | | | | | | |
| 5H | Max Ampl (mPFC) | $mPFC_{CTB+}$ | 11 | 5.49 ± | 1.46 | GLME | maxampl ~ group + (1\|rec)' | | | | |
| | | $mPFC_{CTB-}$ | 10 | 2.79 ± | 0.58 | Term | Fstat | DF1 | DF2 | p | |

| | | | | | | | | | | | | | | | | | |
|---|---|---|---|---|---|---|---|---|---|---|---|---|---|---|---|---|---|
| | | | | | | Intercept | 0.06 | 1 | 15.00 | 0.814 | | | | | | | |
| | | | | | | Group | 6.95 | 1 | 15.00 | **0.019** | | | | | | | |
| | Max Ampl (IC) | $IC_{CTB+}$ | 2 | 1.39 ± | 0.98 | | | | | | | | | | | | |
| | | $IC_{CTB-}$ | 3 | 2.21 ± | 0.98 | | | | | | | | | | | | |
| Fig S4 | | | | | | | | | | | | | | | | | |
| S4E | Mediolateral Peak | mPFC | 3 | 63.75 ± | 4.21 | GLME | var ~ gr*ap + (1\|sample)' | | | | | | | | | | |
| | | IC | 3 | 86.17 ± | 3.82 | Term | Fstat | DF1 | DF2 | p | | | | | | | |
| | | | | | | Intercept | 843.4573 | 1 | 20.00 | **0.000** | | | | | | | |
| | | | | | | gr | 25.7408 | 1 | 20.00 | **0.000** | | | | | | | |
| | | | | | | ap | 1.2429 | 1 | 20.00 | 0.278 | | | | | | | |
| | | | | | | gr:ap | 3.0883 | 1 | 20.00 | 0.094 | | | | | | | |
| S4F | Dorsolateral Peak | mPFC | 3 | 23.42 ± | 4.21 | GLME | var ~ gr*ap + (1\|sample)' | | | | | | | | | | |
| | | IC | 3 | 18.33 ± | 3.99 | Term | Fstat | DF1 | DF2 | p | | | | | | | |
| | | | | | | Intercept | 66.755 | 1 | 20.00 | **0.000** | | | | | | | |
| | | | | | | gr | 5.3966 | 1 | 20.00 | **0.031** | | | | | | | |
| | | | | | | ap | 1.1399 | 1 | 20.00 | 0.298 | | | | | | | |
| | | | | | | gr:ap | 2.54 | 1 | 20.00 | 0.127 | | | | | | | |
| S4G | Total Area Fraction | mPFC | 3 | 0.28 ± | 0.04 | GLME | var ~ gr*ap + (1\|sample)' | | | | | | | | | | |
| | | IC | 3 | 0.12 ± | 0.04 | Term | Fstat | DF1 | DF2 | p | | | | | | | |
| | | | | | | Intercept | 7.7405 | 1 | 20.00 | **0.012** | | | | | | | |
| | | | | | | gr | 57.9087 | 1 | 20.00 | **0.000** | | | | | | | |
| | | | | | | ap | 16.3309 | 1 | 20.00 | **0.001** | | | | | | | |
| | | | | | | gr:ap | 23.72 | 1 | 20.00 | **0.000** | | | | | | | |
| Fig S5 | | | | | | | | | | | | | | | | | |
| S5L | Desensitization | $ChrimsonR_{LED590}$ | 7 | -41.92 ± | 11.69 | 'Source' | 'SS' | 'df' | 'MS' | 'Chi-sq' | p | Wilcoxon Signed rank test | | | zval | p | FDR |
| | | $ChrimsonR_{LED470}$ | 7 | -31.15 ± | 8.78 | 'Columns' | 518.00 | 2 | 259 | 13.77 | 0.001 | ChrimsonR | LED590 | LED470 | -1.521 | 0.128 | 0.128 |
| | | $ChrimsonR_{LED470-D}$ | 7 | -0.08 ± | 0.08 | 'Error' | 234.50 | 18 | 13.02778 | | | ChrimsonR | LED590 | Desentizit | -2.366 | 0.018 | **0.027** |
| | | | | | | 'Total' | 752.50 | 20 | | | | ChrimsonR | LED470 | Desentizit | -2.366 | 0.018 | **0.027** |
| Fig S6 | | | | | | | | | | | | | | | | | |
| S6B | Firing Rate | $mPFC_{Preferring}$ | 15 | | | Repeated Measures ANOVA | | | | | | | | | | | |
| | | $IC_{Preferring}$ | 7 | | | | SumSq | DF | MeanSq | F | pValue | pValueGG | pValueHF | pValueLB | | | |
| | | | | | | (Intercept):Time | 9543.50 | 24 | 397.64 | 2.34 | **0.000** | 0.086 | 0.074 | 0.142 | | | |
| | | | | | | group:Time | 3706.20 | 24 | 154.43 | 0.91 | 0.589 | 0.438 | 0.450 | 0.352 | | | |
| | | | | | | Error(Time) | 81503.00 | 480 | 169.8 | | | | | | | | |
| S6C | Gain | $mPFC_{Preferring}$ | 15 | 2.60 ± | 0.41 | Wilcoxon rank sum | n1 | n2 | zval | p | | | | | | | |
| | | $IC_{Preferring}$ | 7 | 2.36 ± | 0.51 | | 15 | 7 | 0.07 | 0.944 | | | | | | | |
| S6D | Rheobase | $mPFC_{Preferring}$ | 15 | 30.57 ± | 7.42 | Wilcoxon rank sum | n1 | n2 | zval | p | | | | | | | |
| | | $IC_{Preferring}$ | 7 | 36.23 ± | 15.52 | | 15 | 7 | -0.3525 | 0.725 | | | | | | | |
| S6F | sEPSC frequency | $mPFC_{Preferring}$ | 21 | 5.89 ± | 1.42 | Wilcoxon rank sum | n1 | n2 | zval | p | | | | | | | |
| | | $IC_{Preferring}$ | 10 | 5.07 ± | 1.10 | | 21 | 9 | -0.7241 | 0.469 | | | | | | | |
| S6G | sEPSC amplitude (abs) | $mPFC_{Preferring}$ | 21 | 28.39 ± | 5.60 | Wilcoxon rank sum | n1 | n2 | zval | p | | | | | | | |
| | | $IC_{Preferring}$ | 10 | 19.3682 ± | 2.61 | | 21 | 9 | 0.23 | 0.821 | | | | | | | |
| S6H | P (oEPSC > sEPSC) | $mPFC_{Preferring}$ | 21 | 0.51 ± | 0.09 | Wilcoxon rank sum | n1 | n2 | zval | p | | | | | | | |
| | | $IC_{Preferring}$ | 10 | 0.29 ± | 0.12 | | 21 | 9 | 1.61 | 0.108 | | | | | | | |
