## Supplementary Figures and Table S1 for "Integration of cortical inputs in the lateral hypothalamus is dominated by the medial prefrontal cortex"

### Document S1

Figures S1-S6

Table S1

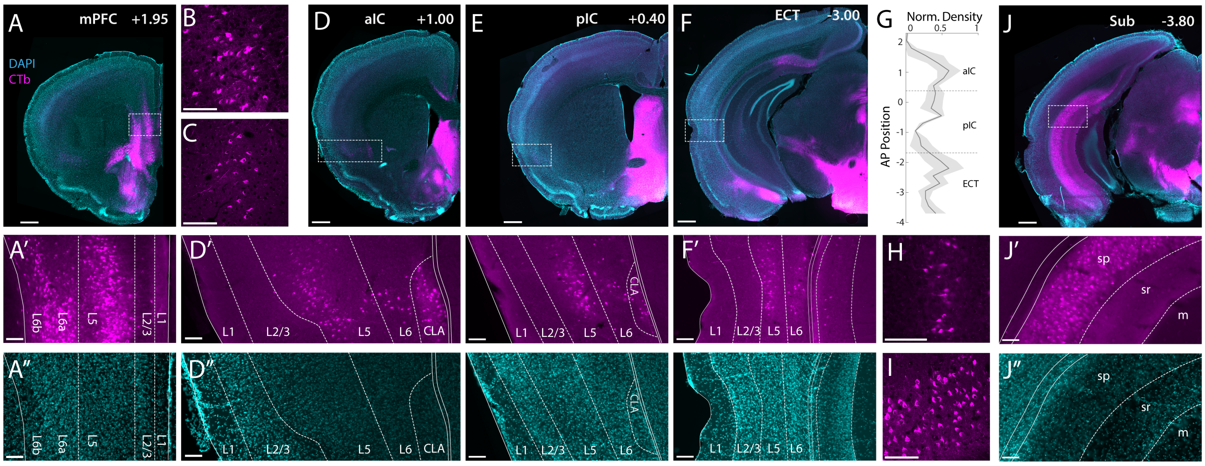


**Figure S1. Distribution of retrograde labelled neurons (Related to Fig. 1)**
(A )Representative section containing the mPFC, AP position is +1.95 from bregma. Scalebar represents 500µm. The bounded box is depicted in A' (CTb) and A" (DAPI) to illustrate the distribution of labelled neurons in layers. Scalebars represent 100µm.
(B) representative confocal image (40x) of labelled pyramidal neurons in L5 of the mPFC .
(C) representative confocal image (40x) of labelled pyramidal neurons in L5 of the aIC.
(D-F) like A, but for aIC (D), pIC (F) and ECT.
(G) Normalized density of labelled neurons in lateral associative areas (aIC, pIC, ECT) for sections ranging from antero-posterior positions from +2 mm to -3.8mm from bregma (n=3 brains).
(H) representative image (20x) of labelled pyramidal neurons in L5 of the ECT.
(I) representative confocal image (40x) of labelled pyramidal neurons in the subiculum.
(J) Similar to ( A & D-F), but for the subiculum (Sub).

Abbreviations: medial prefrontal cortex (mPFC), anterior insular cortex (aIC), claustrum (CLA), posterior insular cortex (pIC), ectorhinal cortex (ECT), subiculum (Sub), pyramidal layer (sp), stratum radiatum (sr), molecular layer (m).

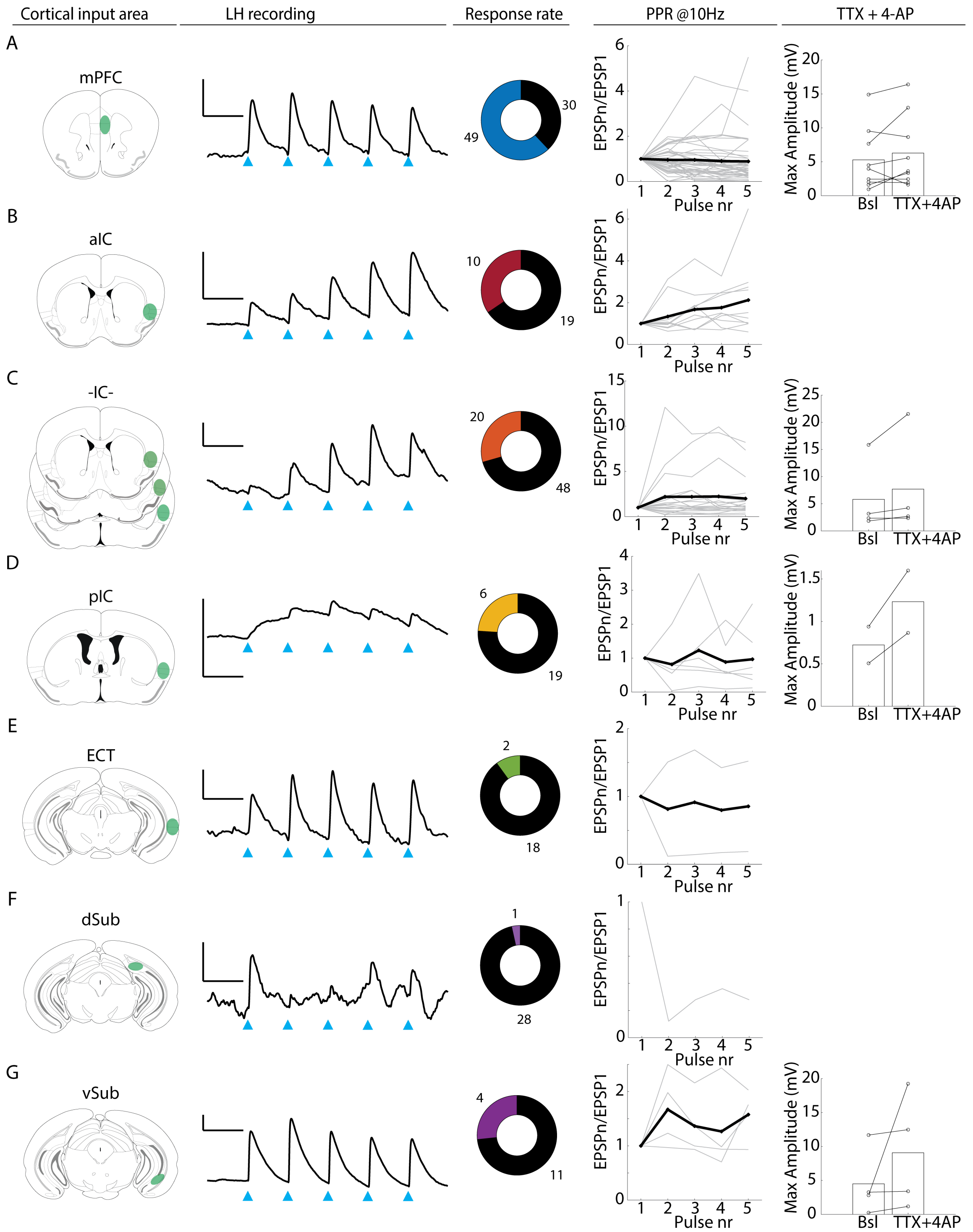


**Supplementary Figure 2. Verification of cortical inputs using channelrhodopsin assisted circuit mapping (Related to Fig. 2)**

For each cortical input area (from left to right): a schematic of the injection area, an example recording of a post-synaptic LH neuron (scalebars are 3mV and 100ms), the response rate, the progressive pulse ratio, and response amplitudes at baseline and after addition of 1µm Tetrodoxin (TTX) and 250 µM 4-aminopyridin (4-AP) . Photostimulation consisted of 5 pulses of 3ms at 10Hz.

Abbreviations: mPFC (medial prefrontal cortex), aIC (anterior insular cortex), -IC- (insular cortex, distributed injection), pIC (posterior insular cortex), ECT (ectorhinal cortex), dSub (dorsal subiculum), vSub (ventral subiculum).

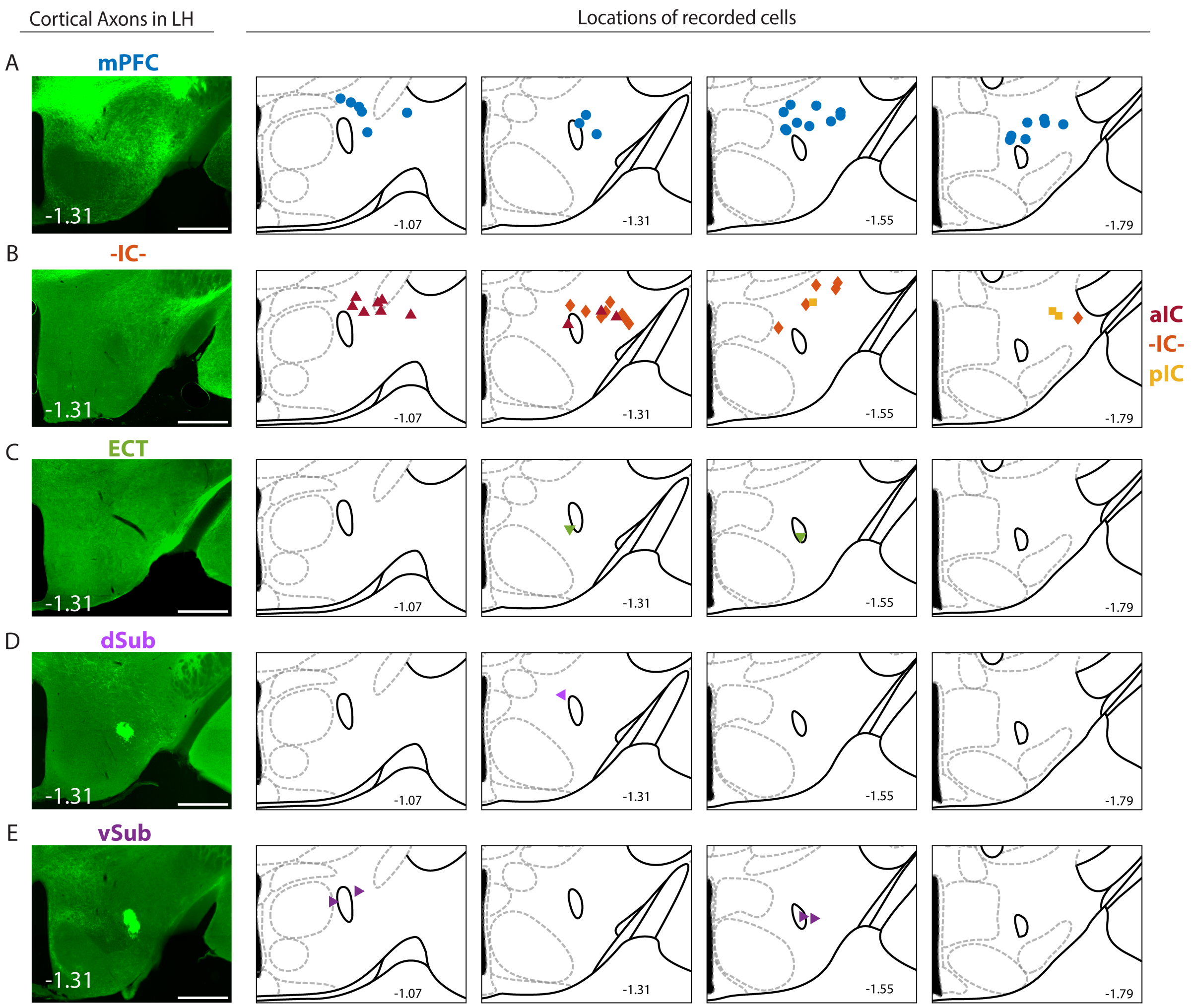


**Figure S3. Topographical organization of cortical axons and inputs (Related to Fig. 2)**

Cortical axons are shown in the representative LH sections (bregma -1.31), scalebars = 500 µm. Locations of responding neurons in LH are depicted as colored shapes, in sections ranging from -1.07 to -1.79 from bregma.

Abbreviations: mPFC (medial prefrontal cortex), aIC (anterior insular cortex), -IC- (insular cortex, distributed injection), pIC (posterior insular cortex), ECT (ectorhinal cortex), dSub (dorsal subiculum), vSub (ventral subiculum).

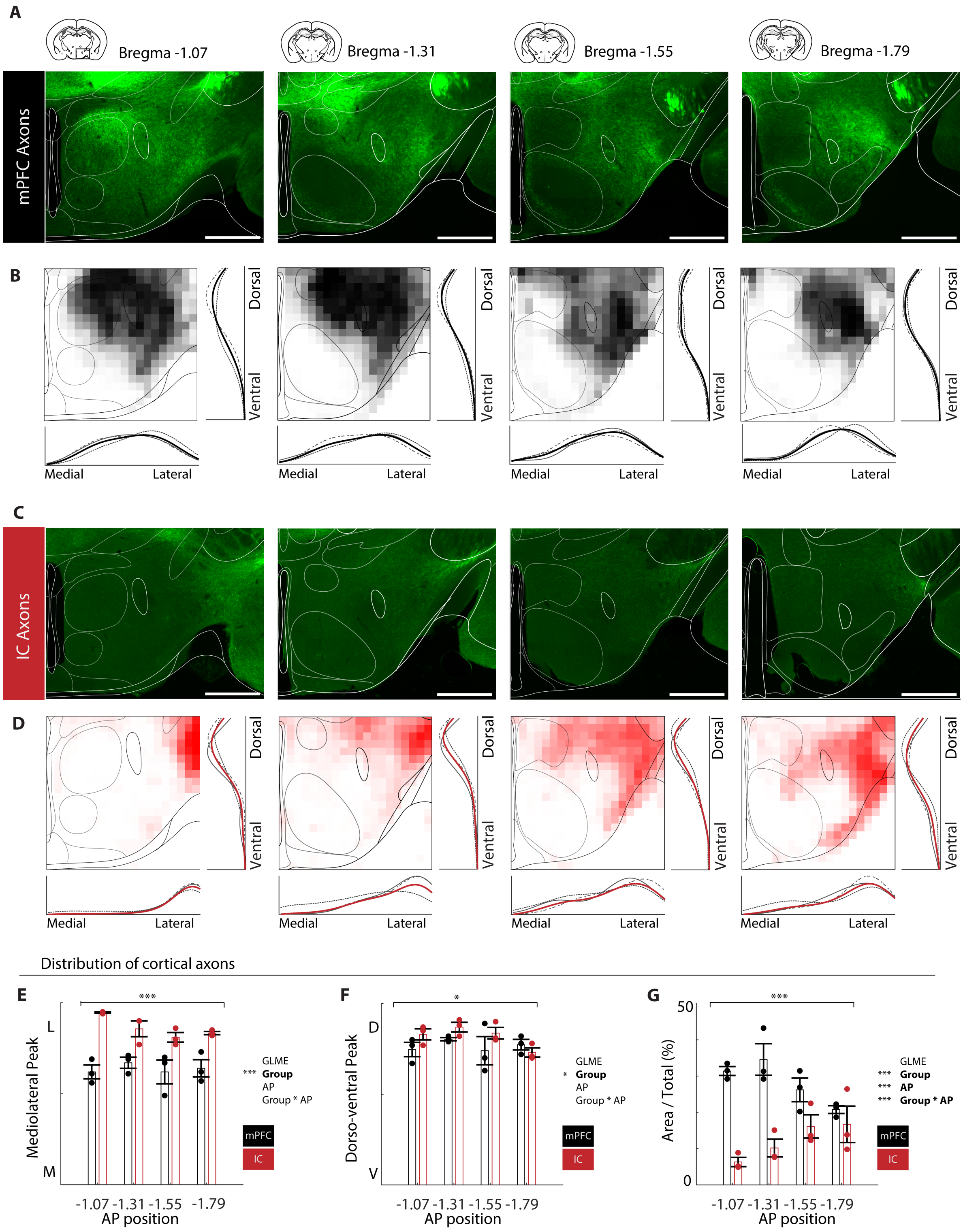


**Figure S4. mPFC and IC target distinct anatomical subdomains (Related to Fig. S3)**

**(A)** Representative examples of mPFC axons in sections containing the LH, in four different anterior posterior (AP) positions.
**(B)** Heatmaps from averaged mPFC axonal densities across 3 different samples, with 1D Ks densities in the mediolateral axis and dorsoventral axis.
**(C-D)** Same as A-B but for IC axons (n=3).

**(E)** Peak densities of mPFC and IC axons in the mediolateral axis, in four different anteroposterior (AP) positions. GLME revealed a significant effect of group (mPFC vs. IC; ($F(1,20) = 25.74$, $p < 0.001$), but not AP position ($F(1,20) = 1.24$ , $p = 0.278$) and their interaction ($F(1,20) = 3.09$, $p = 0.094$).
**(F)** Peak densities of mPFC and IC axons in the dorsoventral axis, in four different anteroposterior (AP) positions. GLME revealed a significant effect of group (mPFC vs. IC; ($F(1, 20) = 5.40$, $p = 0.031$), but not AP position ($F(1, 20) = 1.14$, $p = 0.298$) and their interaction ($F(1, 20) = 2.54$, $p = 0.127$).
**(G)** Proportion of total area covered by mPFC and IC axon, in the four different anterior posterior positions. GLME revealed significant effects of group (mPFC vs. IC; ($F(1,20)=57.91$, $p <0.001$), AP position ($F(1,20)= 16.33$, $p <0.001$) and their interaction ($F(1,20)= 23.72$, $p <0.001$).

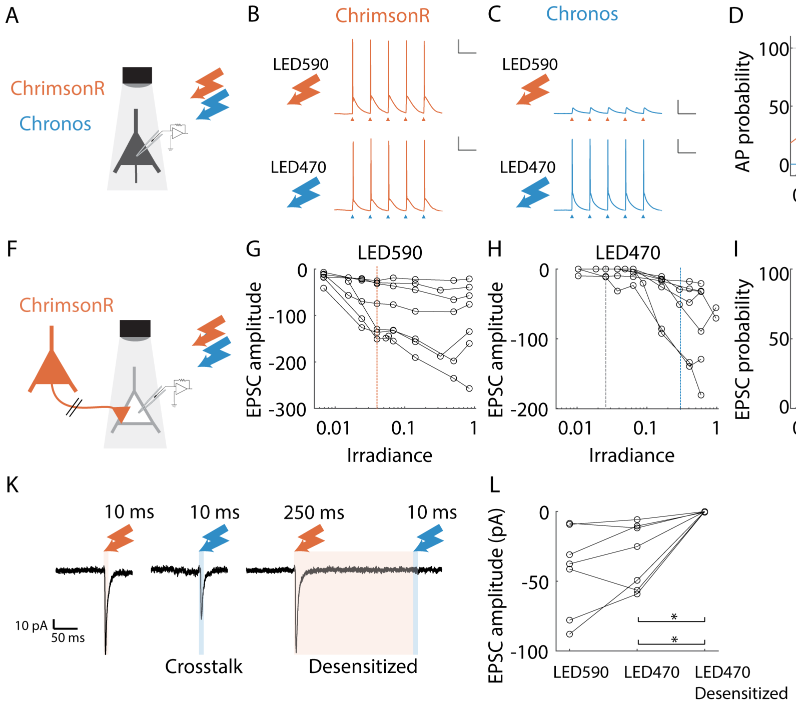


**Figure S5.Verification and optimization of dual channel optogenetics strategy (Related to Fig. 3)**

**(A)** Schematic of photostimulation and recording strategy. ChrimsonR or Chronos expressing neurons at the injection site were recorded to assess light sensitivity to 470nm and 590nm ($N_{Chrimson}$ = 7 cells, $N_{Chronos}$ = 6 cells).
**(B)** Example traces of a Chrimson-expressing neuron ( -68 mV). Light stimulation (5 pulses of 3ms at 10Hz) with 590nm light induced reliable firing with a minimal irradiance of 0.016 mW/mm$^2$ (top panel). In this example, 470nm light induced firing with a minimal irradiance of 0.088 mW/mm$^2$ (bottom panel). Scalebars represent 20mV and 100ms.
**(C)** Same as (B) but for Chronos-expressing neuron ( -65 mV). 590 nm light did not induce firing, even at its maximal power of 1 mW/mm$^2$ (top panel). In this example, 470nm light induced firing with a minimal irradiance of 0.025 mW/mm$^2$ (bottom panel).
**(D)** AP probability in opsin-expressing neurons for different irradiance powers for 590nm wavelength stimulation. The dashed orange line indicates the minimum irradiance needed for all recorded Chrimson neurons to reach a firing probability of 100%. Data are represented as mean ± SEM.
**(E)** Same ad D, but for photostimulation with blue light (470nm). The minimum irradiance needed for reliable spiking at 100% for all recorded Chronos neurons (0.29 mW/mm$^2$) is indicated by a blue dashed line, and for the example in the (C) by a grey line.
**(F)** Schematic of photostimulation and recording strategy. Animals were injected only with AAV-Syn-ChrimsonR-tdT. Recordings were made post-synaptically in LH.
**(G)** EPSC amplitude in response to photostimulation with 590nm light of increasing power (n = 7 cells).
**(H)** Same as G, but for 470nm light.
**(I)** EPSC probability in response to photostimulation with 590nm light of increasing power.
**(J)** Same as I, but for 470nm light.
**(K)** Example traces of a postsynaptic LH neuron, showing that both photo-stimulation of ChrimsonR terminals with 590nm and 470nm wavelength light activates EPSCs. Crosstalk from 470 nm light is suppressed when stimulation is preceded by a long 590nm light pulse, desensitizing ChrimsonR.
**(L)** EPSC amplitudes in response to 10ms stimulation with 590nm light and to 470nm light (crosstalk), and to 470nm light preceded by a desensitizing 590nm light pulse. Crosstalk is suppressed after desensitization (n = 7 cells; $Z_{LED590\ vs.\ LED470}$ = -1.52, p = 0.128; $Z_{LED590\ vs.\ Desensitized\ LED470}$ = -2.37, p = 0.027; $Z_{LED470\ vs.\ Desensitized\ LED470}$ = -2.37, p = 0.027).

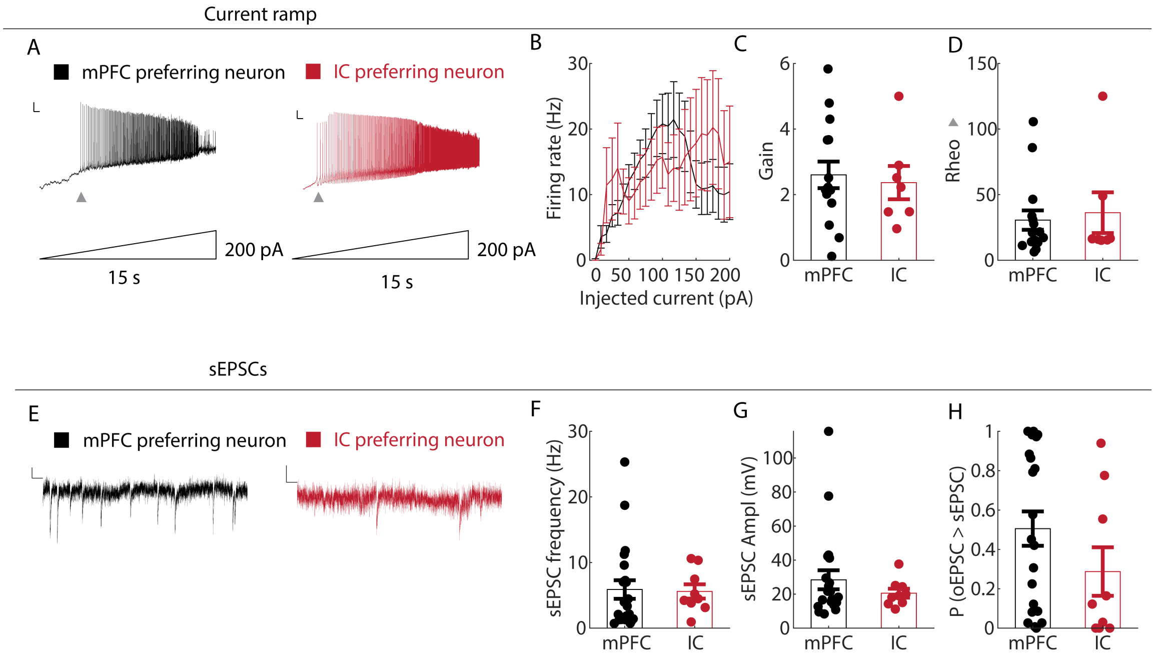


**Figure S6. Firing rate and spontaneous activity of mPFC and IC preferring neurons. (Related to Fig. 4)**

**(A)** Example traces to a current ramp (0-200pA, 15 sec duration). Scalebars represent 10mV and 500ms. Grey triangles the minimal injected current required to start spiking (= Rheo, see H).

**(B)** The firing rate of mPFC- vs. IC- preferring neurons (n=15 vs n=7). Repeated measures ANOVA indicates that firing rate increases with injected current, without any effect of cell type (Ftime (24, 480) = 2.34, p < 0.001; Fgroup*time (24, 480) = 0.91, p = 0.589).

**(C)** The gain, defined as the slope of the firing rate between the first AP and the peak firing rate, was similar for both populations (Z = 0.071, p = 0.944) .

**(D)** The injected current required to start spiking (= Rheo), did not differ between response types (Z = -0.35, p = 0.724).

**(E)** Example traces showing spontaneous EPSCs (sEPSCs) in a mPFC preferring (black trace) and IC preferring neuron (red trace).

**(F)** The frequency of sEPSCs for mPFC- and IC-preferring neurons (Z = -0.72, p = 0.469).

**(G)** The amplitude of sEPSCs was similar for mPFC- and IC-preferring neurons (Z = 0.23, p = 0.821).

**(H)** Plot showing for each neuron the proportion of sEPSCs with amplitudes equal to or smaller than the amplitude of the largest optically evoked EPSC ($EPSC_{mPFC}$ or $EPSC_{IC}$ (Z = 1.61, p = 0.108).

### Tables

| | | % of labelled cells | | | Density | | |
|---|---|---|---|---|---|---|---|
| **Area** | **n** | **Mean** | | **SEM** | **Mean** | | **SEM** |
| PL | 3 | 1.70 | ± | 1.00 | 107.03 | ± | 66.58 |
| IL | 3 | 8.01 | ± | 2.80 | 284.73 | ± | 139.44 |
| ACd | 3 | 0.77 | ± | 0.14 | 26.27 | ± | 8.17 |
| ACv | 3 | 1.37 | ± | 0.37 | 60.81 | ± | 14.41 |
| ORB | 3 | 2.21 | ± | 1.05 | 115.06 | ± | 50.87 |
| IC | 3 | 2.30 | ± | 0.60 | 45.59 | ± | 33.00 |
| CLA | 3 | 2.34 | ± | 0.47 | 208.05 | ± | 116.70 |
| ECT | 3 | 1.75 | ± | 0.80 | 33.74 | ± | 18.04 |
| dSub | 3 | 11.28 | ± | 6.80 | 417.05 | ± | 219.02 |
| CA1 | 3 | 4.13 | ± | 1.38 | 259.62 | ± | 24.91 |
| vSub | 3 | 11.30 | ± | 2.12 | 499.50 | ± | 310.89 |
| PIR | 3 | 1.97 | ± | 0.97 | 10.20 | ± | 9.21 |
| CoA | 3 | 5.42 | ± | 0.33 | 132.76 | ± | 82.55 |
| TT | 3 | 4.37 | ± | 2.08 | 322.47 | ± | 219.90 |
| LS | 3 | 22.53 | ± | 9.27 | 311.66 | ± | 87.08 |
| AMY | 3 | 15.65 | ± | 5.26 | 154.13 | ± | 113.22 |
| NAc | 3 | 2.89 | ± | 1.83 | 64.93 | ± | 70.94 |

**Table S1. Retrograde labelling (Related to figure 1)**
